# A Protein A based *Staphylococcus aureus* vaccine with improved safety

**DOI:** 10.1101/2021.02.19.432007

**Authors:** Miaomiao Shi, Xinhai Chen, Yan Sun, Hwan Keun Kim, Olaf Schneewind, Dominique Missiakas

**Affiliations:** Howard Taylor Ricketts Laboratory, Argonne National Laboratory, Lemont, IL 60439

**Keywords:** *Staphylococcus aureus*, vaccine, protein A, anaphylaxis, IgE, B cell, superantigen, bloodstream infection, colonization, safety

## Abstract

Exposure to *Staphylococcus aureus* does not lead to immunity as evidenced by the persistent colonization of one third of the human population. *S. aureus* immune escape is mediated by factors that preempt complement activation, destroy phagocytes, and modify B and T cell responses. One such factor, Staphylococcal protein A (SpA) encompasses five Immunoglobulin binding domains (IgBDs) that associate with the Fcγ domain to block phagocytosis. IgBDs also associate with the Fab domain of V_H_3-idiotypic IgM which activates B cells with the resulting secretion of antibodies that cannot bind determinants of ***S. aureus***. SpA crosslinking of V_H_3-idiotypic IgG and IgE receptors of mast cells and basophils promotes histamine release and anaphylaxis. Previous work demonstrated the safety, immunogenicity, and protective efficacy of SpA_KKAA,_ a variant partially defective for V_H_3-idiotypic Ig cross-linking, in murine models of ***S. aureus***. Compared to mice (10%), humans produce significantly more V_H_3-idiotypic B cells (50%), prompting a search for safer SpA variants that may be suitably developed as clinical-grade vaccines for efficacy testing in humans. Here, we report the identification of such variants.

## INTRODUCTION

*Staphylococcus aureus* colonizes the human nasopharynx and gastrointestinal tract and is found as a persistent colonizer in approximately one third of the human population [1]. Virtually all humans develop antibodies against some of the molecular determinants of *S. aureus* during childhood [2]. However, these immune responses neither affect *S. aureus* colonization nor protect against invasive disease [2, 3]. Several vaccine candidates have been subjected to clinical efficacy testing, yet their products could not achieve their clinical endpoints [4–6]. Such failure can be explained by the formidable array of immune evasion determinants produced by *S. aureus* in particular Staphylococcal protein A (SpA), a highly conserved secreted protein (Fig. 1a) [7–10]. Secreted SpA is anchored to peptidoglycan by sortase A for even display on the cell surface of the bacterium [11]. Large amounts of peptidoglycan-linked SpA are also released during bacterial replication [12]. Cell wall-anchored SpA binds to the Fcγ-domain of most immunoglobulin subclasses and disables the effector functions of antibodies [10, 13]. Released-SpA crosslinks V_H_3 idiotype B cell receptors (BCRs or IgM) and triggers B cell proliferation and the generation of activated plasmablasts that secrete V_H_3-idiotypic immunoglubulins [7, 8]. These antibodies are constantly produced during colonization and infection but cannot bind to the antigenic determinants of *S. aureus* [9, 10, 14]. SpA encompasses five Immunoglobulin-binding domains (IgBDs) designated E, D, A, B, and C (Fig. 1a) [15, 16]. Each module can bind to the Fcγ-domain of human IgG1, IgG2, and IgG4 and to the variant heavy chains of V_H_3 clonal human IgM, IgG, IgE, IgD and IgA [10, 11].

**Fig. 1.**
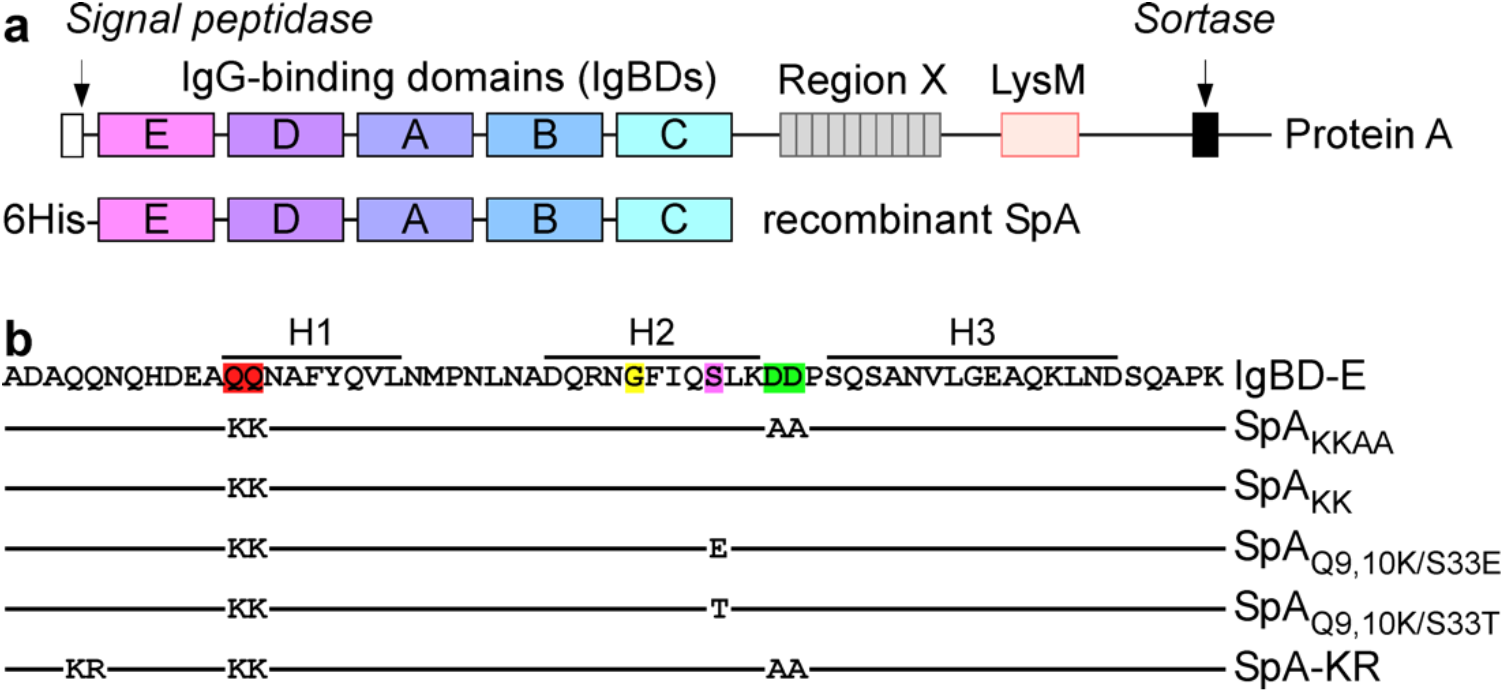
Staphylococcal protein A and vaccine antigens. **a** | The top diagram illustrates the primary structure of Staphylococcal protein A (SpA) with an N-terminal signal peptide cleaved by signal peptidase upon secretion, Immunoglobulin-binding domains (IgBDs), variable Region X made of repeats, LysM domain, and C-terminal LPXTG sorting signal for attachment to peptidoglycan by sortase A. The bottom diagram illustrates the primary structure of recombinant SpA produced in *E. coli* for vaccine studies. **b** | The amino acid sequence of the IgBD-E domain is displayed with positions of three α-helices for each IgBD (H1, H2, and H3). Key amino acids analyzed in this study are indicated in red (Q^9^, Q^10^), yellow (G^29^), pink (S^33^), and green (D^36^, D^37^). These six amino acids are conserved in all five IgBDs. Amino acid numbering is as described by Nilsson *et al*. [26]. Lines under the sequence of IgBD-E depict five SpA variants used in this study with corresponding amino acid substitutions. For simplicity, only one of five IgBD segment is shown.

When injected into mice or guinea pigs, purified SpA does not elicit antibody responses directed against the IgBDs [13, 17]. Similarly, *S. aureus* colonization or invasive disease is not associated with the development of SpA IgBD-specific antibodies in mice, guinea pigs, or humans [7, 9, 10, 17]. The immunoglobulin-binding attributes of SpA are also associated with toxicity. Injection of purified SpA into humans or guinea pigs induces anaphylactic shock, respiratory distress, vascular leakage or death; this can be explained by the abundance of V_H_3-idiotype antibody (50% in humans and 20-30% in guinea pigs)[13, 18, 19]. Mice with only 5-10% of V_H_3-idiotype antibody are resistant to SpA-induced anaphylaxis. However, when treated with intravenous injection of human IgG, mice succumb to SpA injection via histamine release from activated basophils and mast cells [20, 21]. Thus, without modification, SpA is not considered safe for use in humans [19, 22].

Earlier studies demonstrated that substitutions of glutamine residues 9 and 10 (Fig. 1b) with lysine (Q^9,10^K), and of aspartate residues 36 and 37 with alanine (D^36,37^A) disrupted the association of IgBD-D with immunoglobulin [17]. When similar substitutions were introduced into all five IgBDs to generate SpA_KKAA_, a recombinant protein that only encompasses the IgBDs of Staphylococcal protein A appended to an N-terminal six-histidine tag (Fig. 1a), binding activities with human immunoglobulins were greatly reduced [17]. When injected in animals, SpA_KKAA_ elicited high titer serum antibodies that promoted opsonophagocytic killing of *S. aureus* in anti-coagulated mouse, guinea pig and human blood [10, 13, 17] and protected animals against *S. aureus* bloodstream infection [13, 17]. Injection of SpA_KKAA_ in guinea pigs results in profound discomfort. Here, we show that SpA_KKAA_ retains V_H_3-crosslinking activity that accounts for its toxicity and use a systematic approach to identify new vaccine variants with improved safety for future clinical testing.

## RESULTS

### A rational approach for the design of a SpA vaccine

The SpA_KKAA_ antigen with amino acid substitutions Q^9,10^K, D^36,37^A in all IgBDs was generated earlier, guided by three dimensional structures of IgBDs in complex with immunoglobulin. Four hydrogen bonds promote interactions between SpA (B domain numbering, Fig. 1) and Fcγ: Gln^9^ (IgG Ser^254^), Gln^10^ (IgG Gln^311^), Asn^11^ (IgG Asn^434^), and Tyr^14^ (IgG Leu^432^)[15]. Conservation of these residues in all five IgBDs implied a universal mechanism of Fcγ binding that was corroborated experimentally by generating SpA_KKAA_ and SpA_KK_ variants [13, 17]. SpA IgBD residues Gly^29^, Ser^33^, Asp^36^, and Asp^37^ are located along a rim of helix 2 and establish key bonds with V_H_3 framework residues on human immunoglobulin heavy chains [15, 16]. Asp^36,37^Ala amino acid substitutions alone (SpA_AA_) or in combination with Q^9,10^K (SpA_KKAA_) diminished but did not abrogate V_H_3 binding (*vide infra*) [15, 17].

Modified SpA variants have also been developed as ligands for optimal purification of monoclonal antibodies (MAbs)[23, 24]. Most therapeutic MAbs belong to the class of V_H_3-idiotyptic IgG1. Their elution from SpA affinity resin (*e.g.* MabSelect™) requires pH as low as 3.1 causing antibody unfolding and aggregation [25]. The Z domain is a synthetic sequence derived from all five IgBDs with two amino acid substitutions, Ala^1^Val and Gly^29^Ala [24, 26, 27]. The latter substitution removes the alkali-sensitive Asn^28^-Gly^29^ peptide bond [27, 28], allowing for alkali (≥0.1 M NaOH) removal of contaminants during affinity chromatography [23, 25]. The Gly^29^Ala substitution also reduces the binding to the V_H_3 framework of IgG1 and such antibodies can be eluted at pH 3.7 instead of 3.1 [24, 27, 28]. Tandem repeats of two (ZZ), five (ZV) and ten (ZX) Z domains have been developed for purification of MAbs (MabSelectSure™) [26, 29]. Computational calculations using the IgBD-C module suggested that Ser^33^ or Asp^36^ may be critical for thermostability and binding to V_H_3 variant heavy chains [25]. Three variants composed of five IgBD-C domains were generated, each with a single amino acid substitution Gly^29^Ala, Ser^33^Glu or Asp^36^Arg, yielding C-G29A.5d, C-S33A.5d and C-D36A.5d [25]. All three proteins bound to human IgG with an association constant *K*_A_ of 3-5×10^8^ M^−1^ comparable to that of wild-type protein A (*K*_A_ 1.4×10^8^ M^−1^) [25, 30]. When the Fab fragment of trastuzumab, a V_H_3-idiotypic IgG1 MAb, was used as ligand, the affinity of C-G29A.5d was measured as *K*_A_ 4.4×10^5^ M^−1^; C-S33E.5d and C-D36A.5d exhibited nearly a 100-fold reduction in affinity (*K*_A_ 5×10^3^ M^−1^), values that are in agreement with non-specific binding [25]. Informed with this knowledge, we aim to improve the safety of SpA vaccine variants by introducing amino acid substitutions at positions Gly^29^ or Ser^33^ of IgBDs already bearing Q^9,10^K modifications for the disruption of Fcγ interaction [10, 17].

### Amino acid substitutions at Gly^29^ of SpA vaccine candidates

To identify amino acid substitutions at position Gly^29^ of SpA-IgBDs that cause the greatest reduction in affinity between human IgG and SpA_KK_, nineteen plasmids encoding N-terminal six-histidine-tagged SpA_Q9,10K/G29X_, where X is any one of the 19 natural amino acids (except glycine) provided by the genetic code, were constructed. SpA_Q9,10K/G29X_ proteins were purified and bound at equal concentration (250 nM) to Bio-Rad ProteOn HTGchip. Each chip was subjected to SPR experiments with serial dilutions of human IgG or PBS control. The association of human IgG with SpA vaccine candidates loaded on the chip were recorded and data transformed to derive the association constants for each protein (Supplementary Table 1). As a control, the association constants of SpA (*K*_A_ 1.081 × 10^8^ M^−1^) and SpA_KKAA_ for human IgG (*K*_A_ 5.022 × 10^5^ M^−1^) were also quantified. For SpA_Q9,10K/G29X_ proteins, four amino acid substitutions at Gly^29^ caused a significant increase in the association constant: Gly^29^Ser (*K*_A_ 9.398 × 10^5^ M^−1^), Gly^29^Lys (*K*_A_ 9.738 × 10^5^ M^−1^), Gly^29^Ile (*K*_A_ 10.070 × 10^5^ M^−1^), and Gly^29^Ala (*K*_A_ 11.310 × 10^5^ M^−1^), suggesting that these variants bound more tightly to the V_H_3-variant heavy chains of human IgG than SpA_KKAA_ (Supplementary Table 1). The observations for SpA_Q9,10K/G29A_ were surprising to us. The Gly^29^Ala substitution in the ZZZZ construct for commercial antibody purification (MabSelectSure™) diminishes binding to V_H_3-IgG [26], whereas Gly^29^Ala in the context of Gln^9,10^Lys within SpA-IgBDs may promote a modest increase in the affinity for V_H_3-IgG. As compared with SpA_KKAA_, twelve amino acid substitutions at Gly^29^ did not cause a significant difference in the association constant with: Gly^29^Thr, Gly^29^Leu, Gly^29^Glu, Gly^29^Pro, Gly^29^Phe, Gly^29^Met, Gly^29^Val, Gly^29^Trp, Gly^29^Asp, Gly^29^Arg, Gly^29^Asn, and Gly^29^Tyr (Supplementary Table 1). Another three amino acid substitutions at Gly^29^ reduced the association constant: Gly^29^His (*K*_A_ 1.435 × 10^5^ M^−1^), Gly^29^Cys (*K*_A_ 1.743 × 10^5^ M^−1^), and Gly^29^Gln (*K*_A_ 2.057 × 10^5^ M^−1^) to human IgG as compared to SpA_KKAA_ (Supplementary Table 1). Thus, amino acid substitutions at Gly^29^ do not exert a universal effect on the binding of SpA-IgBDs to human IgG. Some amino acid substitutions at Gly^29^ increase the affinity between human IgG and SpA_Q9,10K/G29X_, whereas others are either neutral (exert no significant effect) or diminish the affinity.

### Amino acid substitutions at Ser^33^ of SpA vaccine candidates

To identify amino acid substitutions at position Ser^33^ of the SpA-IgBDs that cause the greatest reduction in affinity between human IgG and SpA_KK_, we constructed nineteen different plasmids encoding N-terminal six-histidine-tagged SpA_Q9,10K/S33X_, where X is any one of the 19 natural amino acids (except serine) provided by the genetic code. All new proteins were purified and subjected to SPR experiments with serial dilutions of human IgG and PBS control to derive association constants (Supplementary Table 2). Two amino acid substitutions at Ser^33^ caused an increase in affinity for human IgG: Ser^33^Gly (*K*_A_ 11.180 × 10^5^ M^−1^) and Ser^33^Ala (*K*_A_ 10.540 × 10^5^ M^−1^), indicating that these variants exhibit greater affinity for human IgG than SpA_KKAA_ (presumably due to increased affinity for V_H_3-variant heavy chains) (Supplementary Table 2). Fourteen amino acid substitutions at Ser^33^ did not cause a significant difference in the association constant: Ser^33^Tyr, Ser^33^Leu, Ser^33^Trp, Ser^33^Val, Ser^33^His, Ser^33^Asn, Ser^33^Met, Ser^33^Arg, Ser^33^Asp, Ser^33^Phe, Ser^33^Gln, Ser^33^Pro, Ser^33^Cys, and Ser^33^Lys (Supplementary Table 2). Three amino acid substitutions at Ser^33^ decreased the affinity for human IgG and SpA_Q9,10K/S33X_: Ser^33^Thr (*K*_A_ 0.386 × 10^5^ M^−1^), Ser^33^Glu (*K*_A_ 0.496 × 10^5^ M^−1^), and Ser^33^Ile (*K*_A_ 1.840 × 10^5^ M^−1^) (Supplementary Table 2). Thus, some amino acid substitutions at Ser^33^ increase the affinity between human IgG and SpA_Q9,10K/S33X_, whereas others are either neutral (exert no significant effect) or diminish the association with human IgG. Of those that diminish the affinity between human IgG, both Ser^33^Glu and Ser^33^Thr exhibit the largest reduction in the association constant (Supplementary Table 2).

### Combining amino acid substitutions at Gly^29^, Ser^33^, and Asp^36,37^ in SpA vaccine candidates

Next, we asked whether combinations of amino acid substitutions at positions Gly^29^, Ser^33^, or Asp^36,37^ of the IgBDs may further alter interactions with human IgG. To address this possibility, three proteins with amino acid substitutions at Ser^33^ were selected because these candidates displayed decreased (SpA_Q9,10K/S33E_,), unaffected (SpA_Q9,10K/S33F_), and increased (SpA_Q9,10K/S33Q_) affinities toward human IgG (Supplementary Table 2). For SpA_Q9,10K/S33E_ (*K*_A_ 0.496 × 10^5^ M^−1^), no additional effect was observed with added substitutions Gly^29^Ala (*K*_A_ 1.265 × 10^5^ M^−1^), Gly^29^Phe (*K*_A_ 1.575 × 10^5^ M^−1^), Asp^36,37^Ala (*K*_A_ 0.568 × 10^5^ M^−1^), Gly^29^Ala/Asp^36,37^Ala (*K*_A_ 1.892 × 10^5^ M^−1^), or Gly^29^Arg (*K*_A_ 4.840 × 10^5^ M^−1^). However, combining Asp^36,37^Ala with either Gly^29^Phe (*K*_A_ 14.850 × 10^5^ M^−1^) or Gly^29^Arg (*K*_A_ 10.240 × 10^5^ M^−1^) increased the affinity of SpA_Q9,10K/S33E_ for human IgG (Supplementary Table 3). When analyzed for SpA_Q9,10K/S33F_ (*K*_A_ 3.902 × 10^5^ M^−1^), whose association constant is not significantly different from that of SpA_KKAA_, we observed similar effects. None of the substitutions altered the affinity of SpA_Q9,10K/S33F_ for human IgG except when Asp^36,37^Ala was combined with either Gly^29^Phe (SpA_Q9,10K/S33Q/D36,37A/Gly29F_ *K*_A_ 12.470 × 10^5^ M^−1^) which led to increased affinity for human IgG as compared to the parent vaccine (Supplementary Table 3). Thus, combining amino acid substitutions at Gly^29^, Ser^33^ and Asp^36,37^ of the SpA-IgBDs does not predictably reduce the affinity for human IgG.

The analysis of vaccine candidates was completed by including two additional variants SpA-KR and SpA_RRVV_ described in the patent applications WO 2015/144653 AI and EP3101027A1, respectively. SpA-KR is a variant of SpA_KKAA_ with two additional amino acid substitutions in the E domain of the IgBD which carries the additional N-terminal amino acid sequence ADAQQN. It was speculated that the two glutamine (QQ) residues may constitute an additional binding site for human IgG and these residues were substituted for KR. SpA_RRVV_ harbors amino acid substitutions at Gln^9,10^ and Asp^36,37^ of each of the five IgBDs of SpA, albeit that the substitutions replace Gln^9,10^ with arginine (R) and Asp^36,37^ with valine (V). When analyzed for their affinity to human IgG, the association constants of SpA-KR (*K*_A_ 5.464 × 10^5^ M^−1^) and SpA_RRVV_ (*K*_A_ 5.609 × 10^5^ M^−1^) were similar to that of SpA_KKAA_ (Supplementary Table 3).

In summary, the association constant with human IgG was measured for sixty-seven new SpA variants (Supplementary Tables 1-3). Two variants, SpA_Q9,10K/S33E_ and SpA_Q9,10K/S33T_, exhibited the largest and most significant reduction in binding (Supplementary Table 2) and were selected as candidate vaccines for additional characterization.

### Binding activity of SpA candidate vaccines to Fcγ and F(ab)_2_ fragments of human IgG

The newly engineered SpA vaccine variants retain the Gln^9,10^Lys amino acid substitutions in their five IgBDs. To confirm that the new substitutions did not regain Fcγ binding activity, Bio-Layer Interferometer was used to determine binding constants to commercially obtained Fcγ fragment (Jackson ImmunoResearch). As expected, SpA-IgBDs exhibited high affinity for Fcγ (*K*_A_ 5.17×10^7^ M^−1^). The Fcγ-binding activity was abolished for SpA_KKAA_ (*K*_A_ 32.68 M^−1^), SpA-KR (*K*_A_ 39.12 M^−1^), SpA_Q9,10K/S33E_ (*K*_A_ 32.68 M^−1^) and SpA_Q9,10K/S33T_ (*K*_A_ 39.91 M^−1^), respectively (Table 1). Thus, the Ser^33^Glu and Ser^33^Thr substitutions do not perturb the effects of the Gln^9,10^Lys on Fcγ-binding in helix 1 of SpA_Q9,10K/S33E_ and SpA_Q9,10K/S33T_.

**Table 1.** The association constant for binding of each combination mutation to Fcγ fragment of human IgG.

|  |  |  |  |
| --- | --- | --- | --- |
| SpA variant<sup>a</sup> | $K_A$ ( M<sup>-1</sup>)<sup>b</sup> | SD<sup>c</sup> | *P* value<sup>d</sup> |
| SpA | 5.17×10<sup>7</sup> | 8.995×10<sup>6</sup> | - |
| SpA<sub>KKAA</sub> | 32.91 | 16.291 | - |
| SpA<sub>Q9,10K/S33E</sub> | 32.68 | 16.414 | ns |
| SpA<sub>Q9,10K/S33T</sub> | 39.91 | 17.081 | ns |
| SpA-KR | 39.12 | 13.348 | ns |
| <sup>a</sup>Test articles were immobilized on Ni-NTA sensor and subjected to Bio-Layer Interferometer (BLI) with increasing concentrations of Fc fragment of human IgG. Data were analyzed from three independent experimental determinations.<br><sup>b</sup>Data were used to derive the association constant ($K_A$) for each test article.<br><sup>c</sup>Data were used to derive the Standard Deviation (SD) for each test article.<br><sup>d</sup>Data were analyzed with One-way ANOVA with Dunnett's Multiple Comparison Test between test article and SpA<sub>KKAA</sub>. Symbols: ns, not significant; *, $P<0.05$; **, $P<0.01$; ***, $P<0.001$; ****, $P<0.0001$. | | | |

To rigorously measure the binding activity of new SpA variants toward V_H_3 idiotypic antibodies, human polyclonal IgG (54% V_H_3 idiotypic variant heavy chains) was cleaved with papain, and F(ab)_2_ fragments were enriched using affinity chromatography on SpA_KK_ [7]. When examined using SPR, the affinity for human V_H_3-F(ab)_2_ enriched preparations, was greatly diminished for SpA_KKAA_ (*K*_A_ 8.27×10^4^ M^−1^), and SpA-KR (*K*_A_ 6.42×10^4^ M^−1^) as compared to SpA (*K*_A_ 1.44×10^7^ M^−1^) (Table 2). Nonetheless both variants retained significant binding activity when compared to SpA_Q9,10K/S33E_ (*K*_A_ 41.24 M^−1^) and SpA_Q9,10K/S33T_ (*K*_A_ 43.55 M^−1^) (Table 1). SpA_Q9,10K/S33E_ and SpA_Q9,10K/S33T_ exhibited binding properties similar to the PBS control (*i.e.* values obtained when no ligand was added). Thus, amino acid substitutions Ser^33^Glu and Ser^33^Thr eliminate V_H_3-IgG binding and represent promising candidates for the selection of variants with reduced V_H_3-Ig crosslinking activities.

**Table 2.** The association constant for binding of each combination mutation to F(ab)_2_ fragment of human IgG.

|  |  |  |  |
| --- | --- | --- | --- |
| **SpA variant<sup>a</sup>** | **$K_A$ ( M<sup>-1</sup>)<sup>b</sup>** | **SD<sup>c</sup>** | ***P* value<sup>d</sup>** |
| SpA | $1.44 \times 10^7$ | $8.193 \times 10^6$ | - |
| SpA<sub>KKAA</sub> | $8.27 \times 10^4$ | $2.76 \times 10^4$ | - |
| SpA-KR | $6.42 \times 10^4$ | $3.80 \times 10^4$ | ns |
| SpA<sub>Q9,10K/S33E</sub> | 41.24 | 5.386 | *** |
| SpA<sub>Q9,10K/S33T</sub> | 43.55 | 5.737 | *** |
| <sup>a</sup>Test articles were immobilized on Bio-Rad ProteOn HTGsensor and subjected to Surface Plasmon Resonance (SPR) with increasing concentrations of F(ab)<sub>2</sub> fragment of human IgG. Data were analyzed from three independent experimental determinations.<br><sup>b</sup>Data were used to derive the association constant ($K_A$) for each test article.<br><sup>c</sup>Data were used to derive Standard Deviation (SD) for each test article.<br><sup>d</sup>Data were analyzed with One-way ANOVA with Dunnett's Multiple Comparison Test between test article and SpA<sub>KKAA</sub>. Symbols: ns, not significant; *, $P < 0.05$; **, $P < 0.01$; ***, $P < 0.001$; ****, $P < 0.0001$. | | | |

### Mouse model for anaphylactic activity of SpA vaccine candidates

Clinical and experimental studies have shown that vascular hyperpermeability is the hallmark of anaphylaxis [31, 32]. Activated mast cells or basophils release vasoactive mediators, including histamine and platelet-activating factor, which induce the anaphylactic response of vascular hyperpermeability by causing vasodilation and endothelial barrier disruption [32]. Intravenous injection of Evans Blue in mice can be used to quantify vascular permeability and because μMT animals do not produce plasma IgG, they can be exploited to model SpA-induced anaphylactic shock [33, 34]. Thus, μMT animals (*n*=4) were injected intradermally with human V_H_3-idiotypic IgG into the ear tissues. 24 hours later, animals were injected intravenously with test article (SpA, variants, or PBS), and next with Evans Blue solution. After 30 min, the dye was extracted from ear tissues for spectrophotometric quantification. Compared with PBS control [34.73 (±) 8.474 ng Evans Blue/mg ear tissue], SpA treatment caused anaphylactic vascular hyperpermeability, releasing 124.9 ng/mg (±26.54 ng/mg) Evans Blue (PBS vs. SpA, *P*<0.0001) (Fig. 2). Administration of SpA_KKAA_ also caused vascular hyperpermeability [70.31 ng/mg (±23.04 ng/mg); PBS vs. SpA_KKAA_, *P*<0.01], albeit at a lower level than SpA (SpA vs SpA_KKAA_, *P*<0.0001) (Fig. 2). In contrast, intravenous administration of 200 μg SpA_Q9,10K/S33E_ [38.57 ng/mg (±15.07 ng/mg); SpA_Q9,10K/S33E_ vs. PBS, not significant] or SpA_Q9,10K/S33T_ [41.43 ng/mg (±13.15 ng/mg); SpA_Q9,10K/S33T_ vs. PBS, not significant] did not elicit vascular hyperpermeability at sites treated with V_H_3-idiotypic human IgG in μMT mice (Fig. 2). The SpA-KR vaccine candidate elicited anaphylactic vascular hyperpermeability similar to that of SpA_KKAA_ (Fig. 2). Thus, unlike SpA and SpA_KKAA_, SpA_Q9,10K/S33E_ and SpA_Q9,10K/S33T_ cannot crosslink V_H_3-idiotypic IgG to promote anaphylactic reactions in μMT mice at sites pretreated with V_H_3-idiotypic human IgG.

**Fig. 2.**
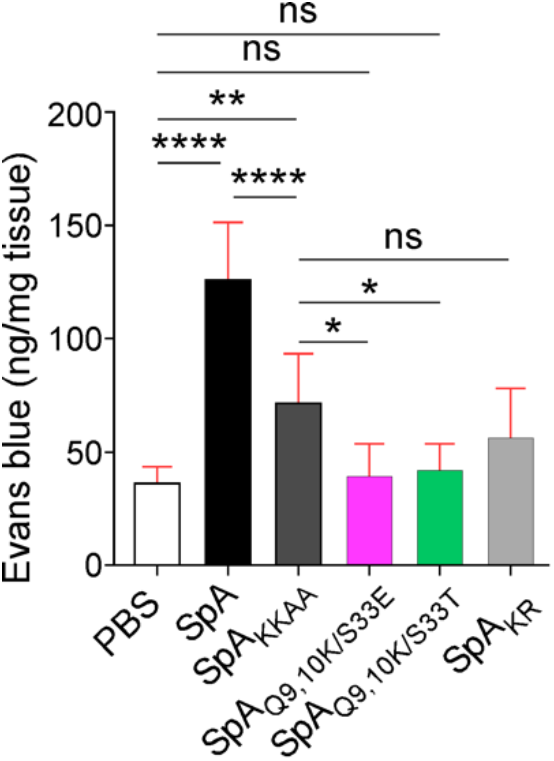
Anaphylactic activity of SpA vaccine candidates in mice. μMT mice (*n*=4) were sensitized with VH3 IgG by intradermal injection in the ear. Candidate vaccine antigens or PBS control were injected intravenously 24 hours later followed by Evans blue injection. After 30 minutes, ear tissues were extracted and extravasation of the dye in these tissues was quantified by spectrophotometric measurement at 620 nm. Data were obtained from three independent experiments and analyzed with ordinary one-way ANOVA with Sidak’s multiple comparison test for statistical differences. Symbols: ns, not significant; *, *P*<0.05; **, *P*<0.01; ***, *P*<0.001; ****, *P*<0.0001.

### SpA vaccine candidate crosslinking of V_H_3-IgE

Basophils and mast cells are two main effector cells of anaphylaxis responses and secrete proinflammatory mediators upon antigen-mediated cross-linking of IgE onto their FcɛRI surface receptors. *S. aureus* Cowan I strain that produces SpA in abundance or soluble purified SpA can activate basophils to induce histamine release. This stimulating effect is dependent on the Fab binding activity of protein A [35]. To study the potential crosslinking effect of SpA vaccine candidates with circulating IgE or IgG bound on the surface of basophils, vaccine variants purified in PBS were added to freshly drawn human blood anti-coagulated with EDTA for 30 min. SpA was used as a positive control. PBS was used as the negative control. Cells were stained with anti-CD123, anti-CD203c, anti-HLA-DR (removal of dendritic cells and monocytes) and anti-CD63. Basophils were identified by gating for SSC^low^CD203c^+^/CD123^+^/HLA-DR^−^ cells (Supplementary Fig. 1). CD123 basophil activation was expressed as a proportion of CD63 and corrected for negative and positive controls. Compared to the PBS control (4.39% activated basophil), SpA or SpA_KKAA_ treatments caused significant increases of CD63^+^ activated basophil population, 32.05% (PBS vs. SpA, *P*<0.0001) and 10.66% (PBS vs. SpA_KKAA_, *P*<0.01), respectively (Table 3). In contrast to SpA_KKAA_, SpA_Q9,10K/S33E_ [5.38%; SpA_Q9,10K/S33T_ vs. SpA_KKAA_, *P*<0.05] or SpA_Q9,10K/S33T_ [4.57%; SpA_Q9,10K/S33T_ vs. SpA_KKAA_, *P*<0.01] were unable to activate basophils and behaved similarly to the PBS control (Table 3). In this assay, SpA-KR [8.15%] and SpA_RRVV_ [10.16%] vaccine candidates showed similar basophil activation as SpA_KKAA_. These data indicate that SpA_Q9,10K/S33E_ and SpA_Q9,10K/S33T_ fail to efficiently crosslink circulating IgE in blood and thus, cannot sensitize basophils by binding the high affinity receptors FcɛRI. The SpA_KKAA_, SpA-KR, and SpA_RRVV_ vaccine candidates appear to retain significant activity for IgE-crosslinking which initiates an unwanted systemic anaphylaxis reaction.

**Table 3.** Activation of human basophils by SpA and vaccine candidate variants.

|  |  |  |  |  |  |  |  |
| --- | --- | --- | --- | --- | --- | --- | --- |
| **SpA variant/**<br>**PBS<sup>a</sup>** | **PBS** | **SpA** | **SpA<sub>KKAA</sub>** | **SpA<sub>Q9,10K/S33E</sub>** | **SpA<sub>Q9,10K/S33T</sub>** | **SpA-KR** | **SpA<sub>RRVV</sub>** |
| % activated basophils | 4.39<br>±0.884 | 32.05<br>±0.919 | 10.66<br>±1.612 | 5.38<br>±0.318 | 4.57<br>±0.877 | 8.15<br>±1.018 | 10.16<br>±0.905 |
| *P* value vs. PBS<sup>b</sup> | - | **** | ** | ns | ns | ns | ** |
| *P* value vs. SpA<sub>KKAA</sub><sup>c</sup> |  | **** | - | * | ** | ns | ns |
| <sup>a</sup>Test articles were incubated with human basophils and data displayed as the percentage of activated basophils of the total basophil population (100%).<br><sup>b,c</sup> One-way ANOVA with Bonferroni's Multiple Comparison Test was performed for statistical analysis that compare test article and PBS<sup>b</sup> or test article and SpA<sub>KKAA</sub><sup>c</sup>. Symbols: ns, not significant; *, *P*<0.05; **, *P*<0.01; ***, *P*<0.001; ****, *P*<0.0001. |  |  |  |  |  |  |  |

Mast cell functional response was measured by antigen-triggered β-hexosaminidase and histamine release. The human mast cell line LAD2 was used for this assay. Mast cells (2×10^5^ cells/ml) were sensitized overnight by incubation with V_H_3 IgE (100 ng/ml) prior to stimulation with SpA vaccine variants (10 μg) for 30 min. Incubation with SpA induced about 35% of β-hexosaminidase release (Fig. 3a). SpA_KKAA_ and SpA-KR vaccines caused 10.32% and 9.87% of β-hexosaminidase release, respectively, with no significant difference (SpA-KR vs. SpA_KKAA_, not significant) (Fig. 3a). These reductions are significant when compared to SpA (SpA vs. SpA_KKAA_, *P*<0.0001; SpA vs. SpA-KR, *P*< 0.0001). Yet, SpA_KKAA_ and SpA-KR vaccines retain β-hexosaminidase releasing activity above negative control levels (SpA_KKAA_ vs. PBS, *P*<0.0001; SpA-KR vs. PBS, *P*< 0.0001) (Fig. 3a). In comparison, SpA_Q9,10K/S33E_ [6.46%; SpA_Q9,10K/S33E_ vs. SpA_KKAA_, *P*< 0.01] and SpA_Q9,10K/S33T_ [4.43%; SpA_Q9,10K/S33T_ vs. SpA_KKAA_, *P*< 0.0001] caused significantly less β-hexosaminidase release as compared to SpA_KKAA_. SpA_Q9,10K/S33E_ and SpA_Q9,10K/S33T_ exhibited similar β-hexosaminidase release as the PBS control (Fig. 3a).

**Fig. 3.**
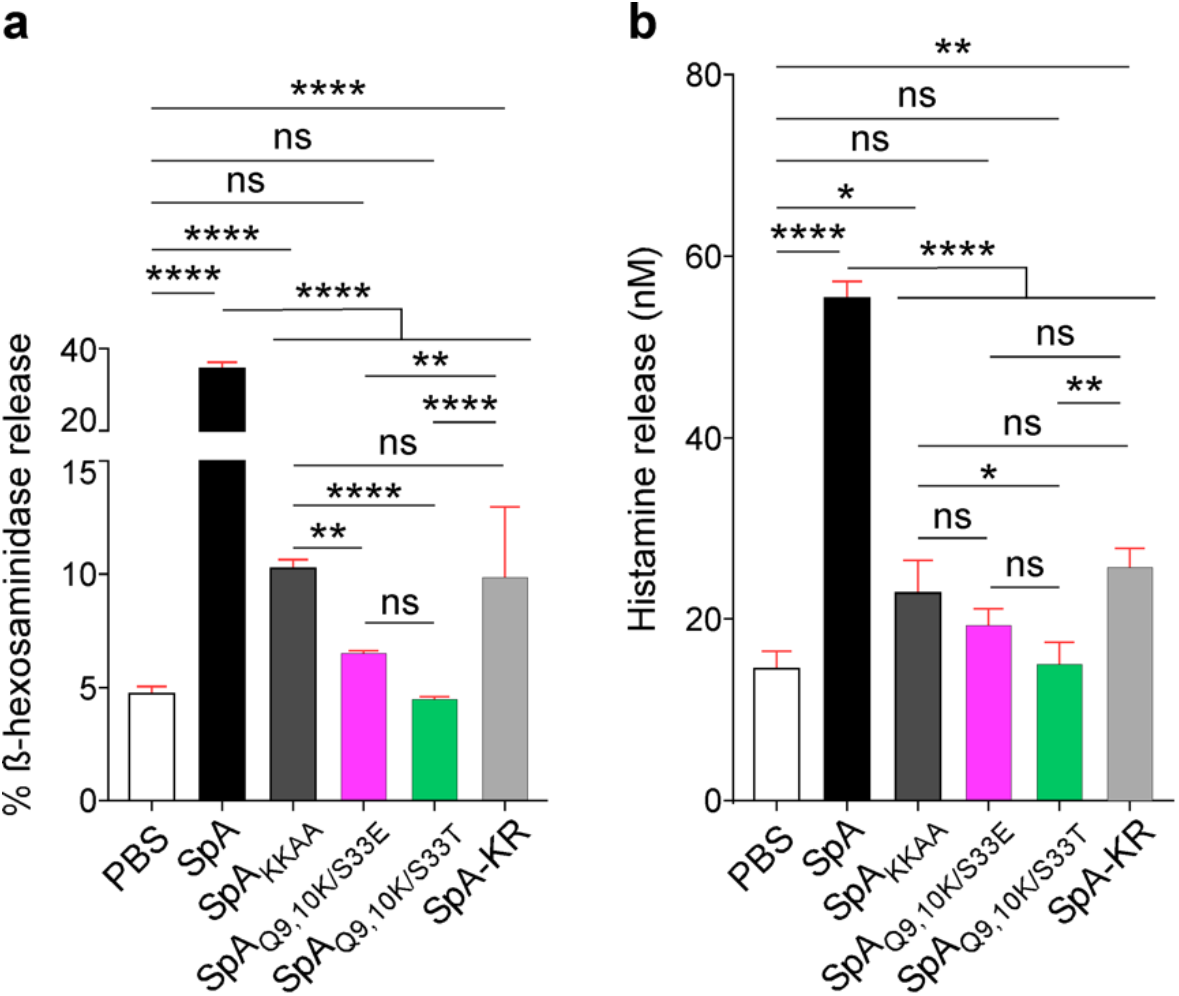
Degranulation of mast cells. Cultured human mast cells (LAD2) were sensitized overnight with V_H_3 IgE, washed, and either left untreated (PBS) or exposed for 1 hour to SpA as a positive control or test articles SpA_KKAA_, SpA_Q9,10K/S33E_, SpA_Q9,10K/S33T_, or SpA-KR. **a** | β-hexosaminidase in the spent medium is shown as the percentage of total β-hexosaminidase released upon treatment of cell cultures with Triton X-100. Measurements were performed in triplicate and one of three experiments is shown. **b** | Amounts of histamine released in the culture medium were measured using an Enzyme Immunoassay. All samples were performed in duplicate. One-way ANOVA with Bonferroni’s multiple comparison test was performed for statistical analysis of the data. Symbols: ns, not significant; *, *P*<0.05; **, *P*<0.01; ***, *P*<0.001; ****, *P*<0.0001.

Similar results were obtained when assessing histamine release (Fig. 3b). SpA stimulated the highest level of histamine release; SpA_KKAA_ and SpA-KR vaccines retained histamine release activity above PBS control levels, and both SpA_Q9,10K/S33E_ and SpA_Q9,10K/S33T_ behaved like the negative control PBS [SpA vs. PBS, or SpA_KKAA_, or SpA-KR, or SpA_Q9,10K/S33E_, or SpA_Q9,10K/S33T_, *P*< 0.0001; SpA_KKAA_ vs SpA-KR or SpA_Q9,10K/S33E_, not significant; SpA_KKAA_ vs. SpA_Q9,10K/S33T_ or PBS, *P*<0.05; SpA_Q9,10K/S33T_ vs. SpA-KR, *P*<0.01].

### Immunogenicity and efficacy of SpA vaccine candidates in the C57BL/6J mouse model of *S. aureus* colonization

*S. aureus* WU1, a member of the multi-locus sequence type ST88 clade, was isolated from an outbreak of preputial gland abscesses among male C57BL/6J mice [9]. Unlike human clinical isolates, *S. aureus* WU1 persistently colonizes the nasopharynx of mice without antibiotic selection and is passed from dams to persistently colonize their offspring [9]. Persistence requires active release of SpA [9]. Immunization of mice with purified SpA_KKAA_ elicits SpA-specific antibodies that induce decolonization of *S. aureus* WU1 from the nasopharynx and gastrointestinal tract of C57BL/6J mice beginning 21 days following intranasal colonization [9]. Decolonization is associated with the generation of antibodies that promote opsonophagocytic killing of *S. aureus* and block the B cell superantigen activity of SpA [9]. To assess the protective attributes of SpA_Q9,10K/S33E_ and SpA_Q9,10K/S33T_ as compared to SpA_KKAA_, cohorts of C57BL/6J mice were immunized with adjuvant alone (mock) or recombinant antigens. Immunization with antigens generated SpA-specific antibodies as measured via ELISA using microtiter plates coated with SpA_KKAA_ or SpA_Q9,10K/S33E_ or SpA_Q9,10K/S33T_, respectively (Fig. 4a). Of note, immunization with SpA_KKAA_ or SpA_Q9,10K/S33T_ elicited antibodies with a bias toward the cognate antigen suggesting that some antibodies may recognize epitopes not shared by all SpA molecules (Fig. 4a). This was not observed upon immunization with SpA_Q9,10K/S33E_. Nonetheless, increases in SpA-antibodies correlated with decolonization of *S. aureus* WU1 from the nasopharynx and gastrointestinal tract of C57BL/6J mice beginning 21 days following intranasal colonization (Fig. 4bc). About 50% of animals cleared bacteria from the nasopharynx and gastrointestinal tract. Animals were bled prior to colonization and at the end of the observation period to derive sera that were used to measure antibody titers against several staphylococcal antigens. IgG levels were significantly increased for several antigens, in particular when animals were immunized with SpA_KKAA_ and SpA_Q9,10K/S33E_ but not with SpA_Q9,10K/S33T_ (Supplementary Table 1). A broader response was observed for animals immunized with SpA_Q9,10K/S33E_ with increased titers observed against ClfB, SdrD, SdrE, SasI, SasD, SasA, FnbpA, FnbpB, SasG, EsxB, SCIN, vWbp and Efb (Supplementary Table 1). Of note, IgG titers against vaccine candidates were sustained over 63 days following colonization.

**Fig. 4.**
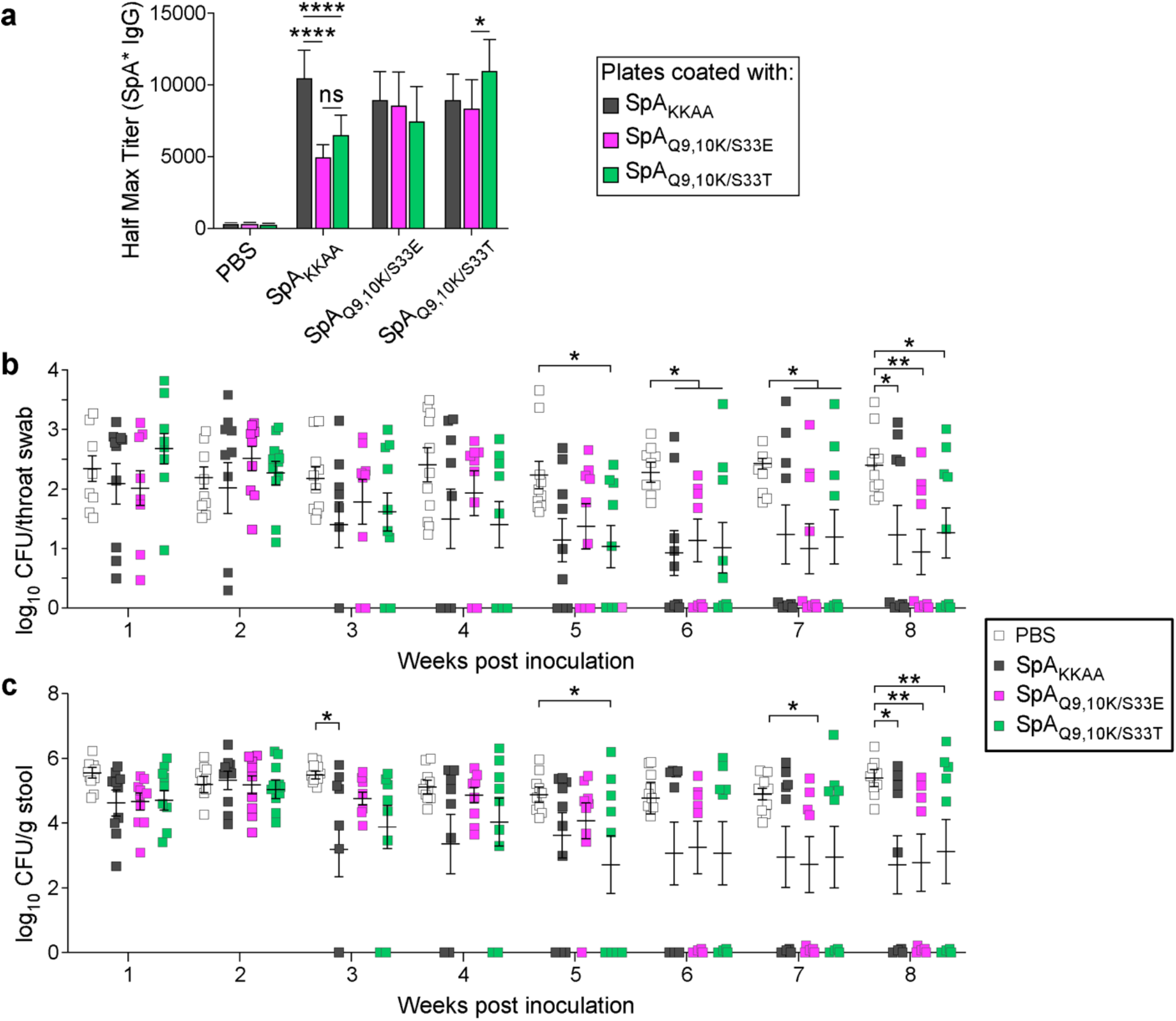
Immunization with SpA_KKAA_ or SpA_Q9,10K/S33E_ or SpA_Q9,10K/S33T_ promotes progressive decolonization. **a** | Three-week-old C57BL/6J mice (*n*=10) were immunized with SpA_KKAA_, SpA_Q9,10K/S33E_, SpA_Q9,10K/S33T_, or PBS control. Mock or booster immunizations occurred on day 11. On day 20, mice were bled to evaluate serum half-maximal antibody titers to vaccine candidates, denoted as SpA* on the *y* axis (**a**). The data was examined with two-way ANOVA with Tukey’s multiple comparisons test. **b-c** | On day 21, animals were inoculated intranasally with 1 × 10^8^ CFU of *S. aureus* WU1. In weekly intervals, mice were swabbed in the throat (**b**) or stool samples were collected (**c**) to enumerate bacterial loads. Each square indicates the number of CFU per throat swab or per gram of stool. The median and standard deviation for each group of animals on a given day are indicated by the horizontal lines and error bars. The data was examined with the two-way ANOVA with Dunnett’s multiple comparison tests (*, *P*<0.05; **, *P*<0.01; ****, *P*<0.0001).

### Immunogenicity and efficacy of SpA vaccine candidates in the BALB/c mouse model of *S. aureus* bloodstream infection

Earlier work demonstrated that immunization of BALB/c mice with SpA_KKAA_ protects animals against intravenous methicillin-resistant *S. aureus* (MRSA) USA300 LAC bloodstream challenge and the ensuing formation of abscess lesions in renal tissues [17]. As compared to mock (adjuvant alone) immunized mice, immunization with SpA_KKAA_, SpA_Q9,10K/S33E_ or SpA_Q9,10K/S33T_ elicited significantly high-titer antibodies against anyone of the three antigens (Fig. 5a). As with C57BL/6J mice, some of the SpA-specific antibody were specific to the antigen used for immunization. This was particularly true for SpA_KKAA_ immunization (Fig. 5a). These results suggest that some (but not all) of the epitopes of antibodies produced by SpA_KKAA_ vaccination are different from that produced by SpA_Q9,10K/S33E_ and SpA_Q9,10K/S33T_ vaccination and *vice versa*. As reported earlier [17], compared to mock-immunized mice, SpA_KKAA_ vaccination reduced the bacterial load of MRSA USA300 LAC and the number of abscess lesions in BALB/c mice (Fig. 5b; *P*<0.0001). SpA_Q9, 10K/S33E_ and SpA_Q9,10K/S33T_ vaccination generated similar protection against MRSA USA300 LAC bloodstream infection compared to SpA_KKAA_ vaccination. Compared to mock-immunized animals, SpA_Q9,10K/S33E_ and SpA_Q9,10K/S33T_ immunization reduced the bacterial load and the number of abscess lesions in BALB/c mice (Fig. 5c; *P*<0.0001). Thus, SpA_Q9,10K/S33E_ and SpA_Q9,10K/S33T_ vaccination elicits similar protection against MRSA USA300 LAC bloodstream infection and associated abscess formation in mice as previously reported for the SpA_KKAA_ vaccine candidate [17].

**Fig. 5.**
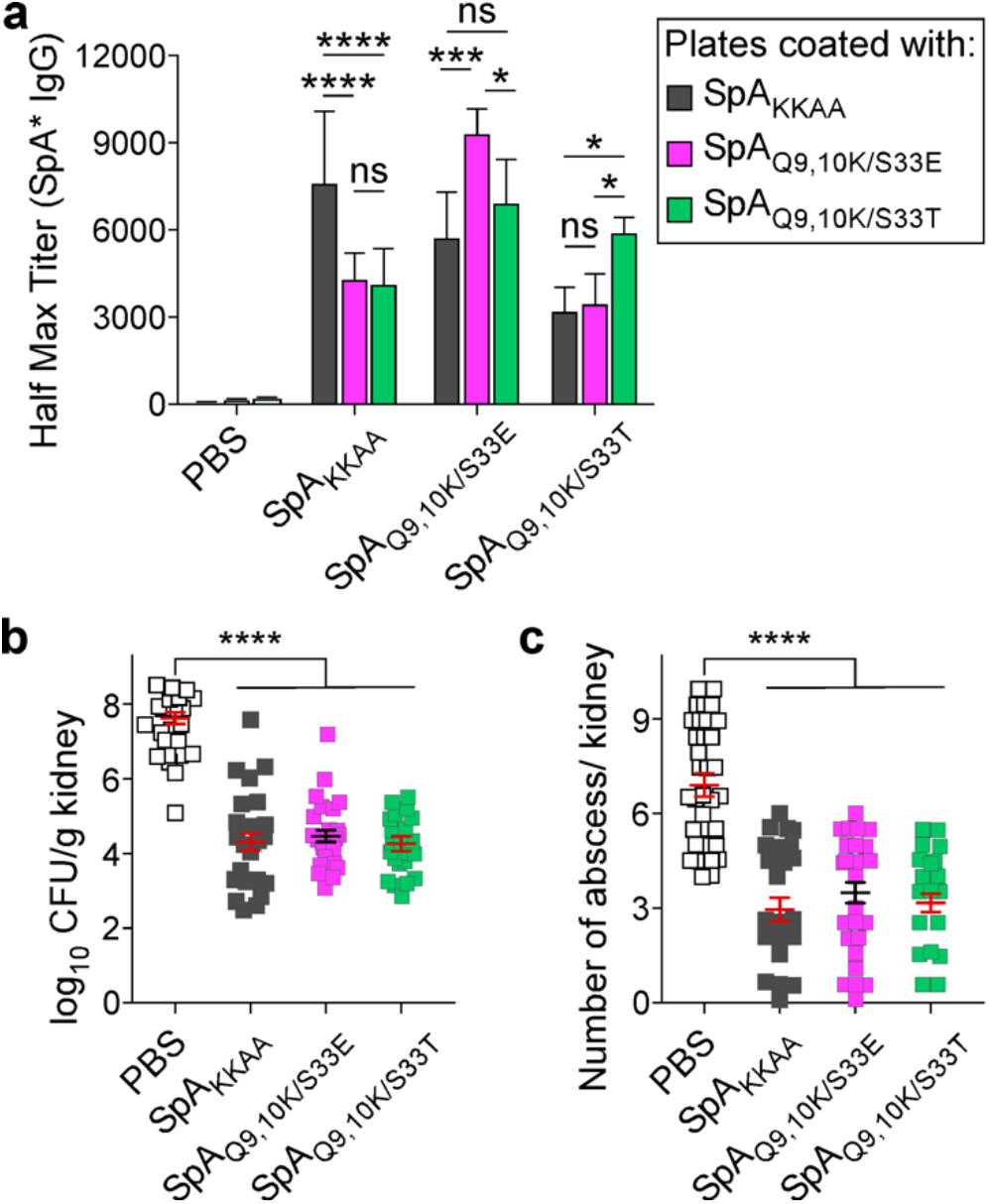
Protective activity of SpA vaccine candidates in the mouse model of bloodstream infection. **a** | Three-week-old BALB/c mice (*n*=15) were immunized as described in Figure 4 and animals were bled to evaluate serum half-maximal antibody titers to vaccine candidates, denoted as SpA* on the *y* axis. On day 21, mice were challenged with 5 × 10^6^ CFU of *S. aureus* USA300 (LAC) into the retroorbital venous sinus of the right eye. **b-c** | Fifteen days post-challenge, animals were euthanized to enumerate staphylococcal loads in kidneys (**b**) and to enumerate abscess lesions (**c**). One-way ANOVA with Bonferroni’s multiple comparison test was performed for statistical analysis of the data (*, *P*<0.05; **, *P*<0.01; ****, *P*<0.0001).

### Binding of SpA vaccine candidates to SpA-neutralizing monoclonal antibody 3F6

Mouse hybridoma monoclonal antibody (hMAb) 3F6 (IgG2a) was derived from SpA_KKAA_-immunized BALB/c mice [36] and used to generate recombinant rMAb 3F6 for production in HEK293 F cells [11]. Both hMAb3F6 and rMAb 3F6 bind to the triple-helical fold of each of the five SpA IgBDs and neutralize their ability to bind human IgG or IgM [11, 36]. Intravenous administration of hMAb3F6 or rMAb 3F6 protects mice against *S. aureus* bloodstream infection [11, 36]. Administration of rMAb 3F6 into mice induces *S. aureus* WU1 decolonization from the nasopharynx and gastrointestinal tract of pre-colonized animals [11]. Here we asked whether rMAb 3F6 binds SpA_Q9,10K/S33E_ or SpA_Q9,10K/S33T_ with similar affinity as SpA_KKAA_ [36]. When measured via ELISA with fixed concentrations of ligands and serial dilutions of rMAb 3F6, affinity constants of 1.51×10^10^ M^−1^, 1.42×10^10^ M^−1^, and 1.34×10^10^ M^−1^ were derived for binding to SpA_KKAA_, SpA_Q9,10K/S33E_, and SpA_Q9,10K/S33T_, respectively (Fig. 6). These data suggest that the amino acid substitutions Ser^33^Glu and Ser^33^Thr do not affect binding of SpA-neutralizing rMAb 3F6. Further, the amino acid substitutions Ser^33^Glu and Ser^33^Thr do not destroy the protective SpA-epitope as defined by the binding of rMAb 3F6.

**Fig. 6.**
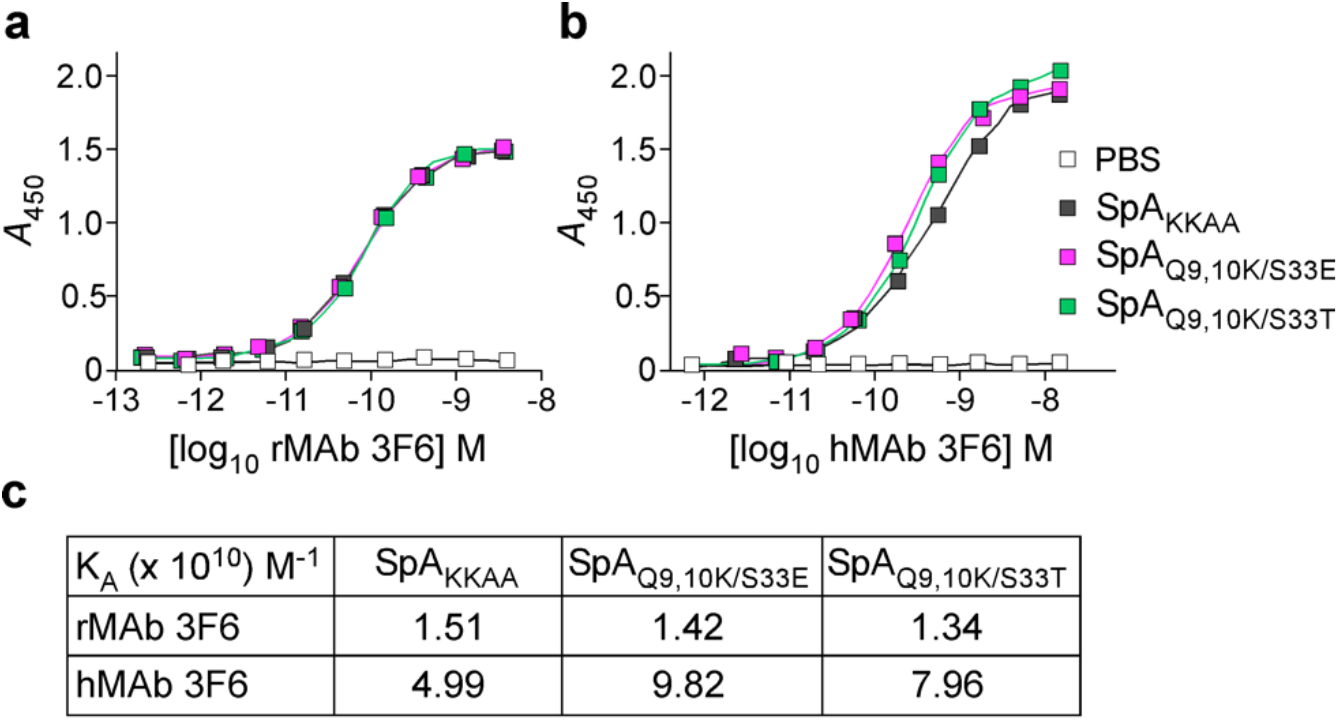
Interaction between SpA vaccine candidates and SpA-neutralizing monoclonal antibody 3F6. **a-b** | Recombinant rMAb 3F6 from HEK293 F cells (**a,**rMAb 3F6) or mouse hybridoma monoclonal antibody (**b**, hMAb 3F6) were serially diluted across enzyme-linked immunosorbent assay plates coated with either SpA_KKAA_, SpA_Q9,10K/S33E_, SpA_Q9,10K/S33T_, or PBS control. **c** | Association constants calculated using GraphPad Prism software.

## DISCUSSION

*S. aureus* causes community- and hospital-acquired invasive diseases, including soft tissue, wound, lung, skeletal, bloodstream and surgical site infections [1, 37]. In 2017, nearly 120,000 *S. aureus* bloodstream infections and 20,000 associated deaths occurred in the United States [38]. While the rate of decline of MRSA bloodstream infections has slowed, bloodstream infections caused by methicillin-susceptible *S. aureus* (MSSA) are increasing slightly in the community (3.9% annually, 2012–2017) [38]. Recurrence is a hallmark of *S. aureus* disease and exemplifies the inability of infected hosts to establish protective immunity [3]. The development of a staphylococcal vaccine that can block colonization and prevent invasive disease has thus far not been achieved.

Vaccines initially evolved from the observation that pre-exposure to some diseases or survival could confer life-long protection. The identification of microbes as causative agents of disease and the ability to isolate and grow such microbes guided earlier efforts for vaccine development. Initially, whole-cell preparations and empirical methods were used to develop vaccines with reduced unwanted side effects [3]. For anthrax disease, culture passaging resulted in heritable attenuation of *Bacillus anthracis* and live attenuated strains have been used extensively to protect animals and humans against anthrax disease [3]. Killed whole-cell preparations were injected in humans for protection against whooping cough caused by *Bordetella pertussis*. However, when whole-cell preparations of *S. aureus* were injected in humans, protection was not observed [39].

Many bacterial pathogens elaborate capsular polysaccharides (capsules) to escape phagocytosis and colonize their host. Chemical conjugation of polysaccharide to protein helps overcome poor immunogenicity [40]. Such conjugate vaccines have been developed successfully to prevent nasopharyngeal colonization and infection by *Haemophilus influenzae* type b, *Neisseria meningitidis*, *Streptococcus pneumoniae*, and *Salmonella typhi* [3]. Other bacterial infections can be explained by the secretion of key toxins. For example, the pathogenesis of diphtheria is critically dependent on diphtheria toxin [41]. Formalin inactivation of diphtheria toxin elicits neutralizing antibodies that protect against the disease when injected in humans [42].

*S. aureus* secretes several toxins and most clinical strains encode genes for either one of two types of capsular polysaccharide [43, 44]. Large efforts were expanded to develop capsular-conjugate vaccines, but efficacy trials revealed that capsule-specific antibodies could not protect against *S. aureus* bacteremia [4, 45]. This failure can be rationalized with the knowledge that not all strains of *S. aureus* produce capsule and thus the capsule is not absolutely required for colonization and infection [44]. However, all strains of *S. aureus* agglutinate, and all strains of *S. aureus* produce Staphylococcal protein A [43, 46]. Agglutination is caused by two secreted coagulases that activate prothrombin nonproteolytically to generate fibrin cables. Fibrin(ogen) binding proteins such as ClfA tether fibrin cables to the bacterial cell surface [46]. Agglutination protects staphylococci from opsonophagocytic killing, and facilitates bacterial dissemination and shielding within abscess lesions [46]. Thus, agglutination is the encapsulation of bacterial cells with host molecules. Another factor, Staphylococcal protein A captures antibodies in non-immune complexes on the cell surface providing a second layer of protection against phagocytes.

We demonstrated earlier that animals immunized with SpA_KKAA_, a vaccine candidate encompassing only the five IgBD of Staphylococcal protein A with four amino acid substitutions in each IgBD, elicited the generation of protective anti-SpA antibodies. However, SpA_KKAA_, as well as a variant SpA-KR, with two additional substitutions, retain significant binding to V_H_3-idiotypic immunoglobulin when using F(ab)_2_ fragment of human IgG as ligand. When analyzed with human basophils (LAD2 cells) coated with V_H_3-IgG, SpA_KKAA_ and SpA-KR trigger V_H_3-Ig crosslinking, as measured by the release of β-hexosaminidase and histamine [35]. In a mouse model for anaphylactic vascular hyperpermeability, the biological effects of such histamine release were measurable as Evans Blue dye extravasation at anatomical sites of V_H_3-IgG administration in μMT mice. Together these observations raise concerns about the safety of SpA vaccine candidates as potential activators of anaphylactic reactions in humans. To address the concerns, we engineered two new antigens, SpA_Q9,10K/S33E_ and SpA_Q9,10K/S33T_, with improved safety profiles. SpA_Q9,10K/S33E_ and SpA_Q9,10K/S33T_ lack affinity for V_H_3-idiotypic immunoglobulins, show reduced or no activity toward histamine release from V_H_3-IgE coated human mast cells, and do not promote Evans Blue dye extravasation in response to V_H_3-IgG injection in μMT mice. Immunization of BALB/c mice with SpA_Q9,10K/S33E_ and SpA_Q9,10K/S33T_ elicited similar levels of SpA-specific IgG responses as SpA_KKAA_. When analyzed for vaccine efficacy in mouse models, vaccination with SpA_Q9,10K/S33E_ or SpA_Q9,10K/S33T_ afforded similar levels of protection against *S. aureus* colonization or invasive bloodstream infection as the SpA_KKAA_ vaccine [17]. Further, the amino acid substitutions Ser^33^Glu and Ser^33^Thr do not perturb the protective IgBD epitopes that are defined by the *S. aureus* colonization- and invasive disease-protective monoclonal antibody 3F6 [11, 36]. Based on these observations, we propose that the *S. aureus* vaccine candidates SpA_Q9,10K/S33E_ and SpA_Q9,10K/S33T_ may be suitable for development as clinical grade vaccines for clinical safety and efficacy testing against *S. aureus* colonization and invasive disease.

## METHODS

Detailed methods are provided in the Supplementary section.

### Ethics statement

The University of Chicago’s Institutional Review Board (IRB) reviewed, approved, and supervised the protocols used for all experiments utilizing blood from human volunteers and informed consent forms were obtained from all participants. Experiments with *S. aureus* were performed in biosafety level 2 (BSL2) and animal BSL2 (ABSL2) containment in accordance with the institutional guidelines following experimental protocol review and approval by the Institutional Biosafety Committee (IBC) and the Institutional Animal Care and Use Committee (IACUC) at the University of Chicago.

### Statistical analyses

For association constant experiments, data were analyzed for significant differences using ordinary one-way ANOVA with Dunnett’s multiple comparisons test between each test article and SpA_KKAA_. For basophil activation experiments, data were analyzed using one-way ANOVA with Bonferroni’s multiple comparisons. For experiments shown in Figures 2 and 3, data were analyzed using ordinary one-way ANOVA with Sidak’s and Bonferroni’s multiple comparisons test, respectively. For experiments shown in Figures 4 and 5, data were analyzed with two-way ANOVA with Tukey’s (Fig. 4a–5a) and Dunnett’s (Fig. 4bc) multiple comparisons tests and, with one-way ANOVA with Bonferroni’s (Fig. 5bc) multiple comparisons test. Serum IgG responses to immunization and colonization (Supplementary Table 4) were analyzed with one-way ANOVA with Sidak’s multiple comparisons test. All data were analyzed with Prism (GraphPad Software, Inc.), and *P* values less than 0.05 were deemed significant.

## Data availability

All data needed to draw the conclusions in the paper are present in the paper and the supplementary material. Additional data related to this paper may be requested from the authors.

## Acknowledgements

We thank Vilasack Thammavongsa, and members of our laboratory for discussion. This project was supported by funds from the National Institute of Allergy and Infectious Diseases, National Institutes of Health, Department of Health and Human Services, under award AI52474 and from the University of Chicago Polsky Center, Innovation Fund.

## REFERENCES

[1] Kluytmans J, van Belkum A, Verburgh H. Nasal carriage of *Staphylococcus aureus*: epidemiology, underlying mechanisms, and associated risks. Clin Microbiol Rev. 1997;10:505–20.

[2] Prevaes SM, van Wamel WJ, de Vogel CP, Veenhoven RH, van Gils EJ, van Belkum A, et al. Nasopharyngeal colonization elicits antibody responses to staphylococcal and pneumococcal proteins that are not associated with a reduced risk of subsequent carriage. Infect Immun. 2012;80:2186–93.

[3] Missiakas D, Schneewind O. *Staphylococcus aureus* vaccines: deviating from the carol. J Exp Med 2016;231:1645–53.

[4] Shinefield H, Black S, Fattom A, Horwith G, Rasgon S, Ordonez J, et al. Use of a *Staphylococcus aureus* conjugate vaccine in patients receiving hemodialysis. N Engl J Med. 2002;346:491–6.

[5] Fowler VG, Allen KB, Moreira ED, Moustafa M, Isgro F, Boucher HW, et al. Effect of an investigational vaccine for preventing *Staphylococcus aureus* infections after cardiothoracic surgery: a randomized trial. JAMA. 2013;309:1368–78.

[6] Harrison KJ. Clinical trial of coagulase and alpha-hemolysin toxoids in chronic furunculosis. Br Med J. 1963;2:149–52.

[7] Pauli NT, Kim HK, Falugi F, Huang M, Dulac J, Dunand CH, et al. *Staphylococcus aureus* infection induces protein A-mediated immune evasion in humans. J Exp Med. 2014;211:2331–9.

[8] Kim HK, Falugi F, Missiakas D, Schneewind O. Peptidoglycan-linked protein A promotes T-cell dependent antibody expansion during *Staphylococcus aureus* infection. Proc Natl Acad Sci USA. 2016;113:5718–23.

[9] Sun Y, Emolo CE, Holtfreter S, Wiles S, Kreiswirth B, Missiakas D, et al. Staphylococcal protein A is required for persistent colonization of mice with *Staphylococcus aureus*. J Bacteriol. 2018;200:e00735–17.

[10] Falugi F, Kim HK, Missiakas DM, Schneewind O. The role of protein A in the evasion of host adaptive immune responses by *Staphylococcus aureus* mBio. 2013;4:e00575–13.

[11] Chen X, Sun Y, Missiakas D, Schneewind O. Staphylococcus aureus Decolonization of Mice With Monoclonal Antibody Neutralizing Protein A. J Infect Dis. 2019;219:884–8.

[12] Becker S, Frankel MB, Schneewind O, Missiakas DM. Release of protein A from the cell wall envelope of *Staphylococcus aureus*. Proc Natl Acad Sci USA. 2014;111:1574–9.

[13] Kim HK, Falugi F, Thomer L, Missiakas DM, Schneewind O. Protein A suppresses immune responses during *Staphylococcus aureus* bloodstream infection in guinea pigs. mBio. 2015;6:e02369–14.

[14] Forsgren A, Svedjelund A, Wigzell H. Lymphocyte stimulation by protein A of *Staphylococcus aureus*. Eur J Immunol. 1976;6:207–13.

[15] Deisenhofer J. Crystallographic refinement and atomic models of a human Fc fragment and its complex with fragment B of protein A from *Staphylococcus aureus* at 2.9- and 2.8-A resolution. Biochemistry. 1981;20:2361–70.

[16] Graille M, Stura EA, Corper AL, Sutton BJ, Taussig MJ, Charbonnier JB, et al. Crystal structure of a *Staphylococcus aureus* protein A domain complexed with the Fab fragment of a human IgM antibody: structural basis for recognition of B-cell receptors and superantigen activity. Proc Nat Acad Sci USA. 2000;97:5399–404.

[17] Kim HK, Cheng AG, Kim H-Y, Missiakas DM, Schneewind O. Non-toxigenic protein A vaccine for methicillin-resistant *Staphylococcus aureus* infections. J Exp Med. 2010;207:1863–70.

[18] Gutafson GT, Stalenheim G, Forsgren A, Sjöquist J. Protein A from *Staphylococcus aureus* IV. Production of anaphylaxis-like cutaneous and systemic reactions in non-immunized guinea pigs. J Immunol. 1968;100:530–4.

[19] Ballow C, Leh A, Slentz-Kesler K, Yan J, Haughey D, Bernton E. Safety, pharmacokinetic, immunogenicity, and pharmacodynamic responses in healthy volunteers following a singlel intravenous injection of purififed staphylococcal protein A. J Clin Pharmacol. 2013;53:909–18.

[20] Gustafson GT, Sjöquist J, Stålenheim G. Protein A from *Staphylococcus aureus*. II. Arthus-like reaction produced in rabbits by interaction of protein A and human gamma-globulin. J Immunol. 1967;98:1178–81.

[21] Anderson AL, Sporici R, Lambris J, Larosa D, Levinson AI. Pathogenesis of B-cell superantigen-induced immune complex-mediated inflammation. Infect Immun. 2006;74:1196–203.

[22] Goldwater R, Garner R, Zamfirov V, Haughey D, Bernton E. PK/PDrelationships in a sequential, escalating, single-dose study of PRTX-100, a highly purified staphylococcal protein A. J Clin Pharmacol. 2007;47:1204.

[23] Minakuchi K, Murata D, Okubo Y, Nakano Y, Yoshida S. Remarkable alkaline stability of an engineered protein A as immunoglobulin affinity ligand: C domain having only one amino acid substitution. Protein Sci. 2013;22:1230–8.

[24] Jansson B, Uhlén M, Nygren PA. All individual domains of staphylococcal protein A show Fab binding. FEMS Immunol Med Microbiol. 1998;20:69–78.

[25] Yoshida S, Murata D, Taira S, Iguchi K, Takano M, Nakano Y, et al. Rational design and engineering of protein A to obtain the controlled elution profile in monoclonal antibody purification. Chem-Bio Informatics Journal 2012;12:1–13.

[26] Nilsson B, Moks T, Jansson B, Abrahmsén L, Elmblad A, Holmgren E, et al. A synthetic IgG-binding domain based on staphylococcal protein A. Protein Eng. 1987;1:107–13.

[27] Gulich S, Uhlen M, Hober S. Protein engineering of an IgG-binding domain allows milder elution conditions during affinity chromatography. J Biotechnol. 2000;76:233–44.

[28] Ghose S, Allen M, Hubbard B, Brooks CP, Cramer SM. Antibody variable region interactions with proteinA: implications for the development of generic purification processes. Biotechnol Bioengin. 2005;92:665–73.

[29] Ljungberg UK, Jansson B, Niss U, Nilsson R, Sandberg BE, Nilsson B. The interaction between different domains of staphylococcal protein A and human polyclonal IgG, IgA, IgM and F(ab’)2: separation of affinity from specificity. Mol Immunol. 1993;30:1279–85.

[30] Svensson HG, Hoogenboom HR, Sjobring U. Protein LA, a novel hybrid protein with unique single-chain Fv antibody and Fab-binding properties. Eur J Biochem. 1998;258:890–6.

[31] Fisher MM. Clinical observations on the pathophysiology and treatment of anaphylactic cardiovascular collapse. Anaesth Intensive Care. 1986;14:17–21.

[32] Korhonen H, Fisslthaler B, Moers A, Wirth A, Habermehl D, Wieland T, et al. Anaphylactic shock depends on endothelial Gq/G11. J Exp Med. 2009;206:411–20.

[33] Nakamura Y, Oscherwitz J, Cease KB, Munoz-Planillo R, Hasegawa M, McGavin MJ, et al. *Staphylococcus* δ-toxin promotes allergic skin disease by inducing mass cell degranulation. Nature. 2013;503 review:397–401.

[34] Kitamura D, Roes J, Kühn R, Rajewsky K. A B cell-deficient mouse by targeted disruption of the membrane exon of the immunoglobulin mu chain gene. Nature. 1991;350:423–6.

[35] Marone G, Tamburini M, Giudizi MG, Biagiotti R, Almerigogna F, Romagnani S. Mechanism of activation of human basophils by *Staphylococcus aureus* Cowan 1. Infect Immun. 1987;55:803–9.

[36] Kim HK, Emolo C, DeDent AC, Falugi F, Missiakas DM, Schneewind O. Protein A-specific monoclonal antibodies and the prevention of *Staphylococcus aureus* disease in mice. Infect Immun. 2012;80:3460–70.

[37] Lowy FD. *Staphylococcus aureus* infections. New Engl J Med. 1998;339:520–32.

[38] Kourtis AP, Hatfield K, Baggs J, Mu Y, See I, Epson E, et al. Vital Signs: Epidemiology and Recent Trends in Methicillin-Resistant and in Methicillin-Susceptible Staphylococcus aureus Bloodstream Infections - United States. MMWR Morb Mortal Wkly Rep. 2019;68:214–9.

[39] Rogers DE, Melly MA. Speculation on the immunology of staphylococcal infections. Ann NY Acad Sci. 1965;128:274–84.

[40] Pollard AJ, Perrett KP, Beverley PC. Maintaining protection against invasive bacteria with protein-polysaccharide conjugate vaccines. Nat Rev Immunol. 2009;9:213–20.

[41] Collier RJ. Understanding the mode of action of diphtheria toxin: a perspective on progress during the 20th century. Toxicon. 2001;39:1793–803.

[42] Glenny AT, Buttle GAH, Stevens MF. Rate of disappearance of diphtheria toxoid injected into rabbits and guinea-pigs: toxoid precipitated with alum. J Pathol Bacteriol. 1931;34:267–87.

[43] Thammavongsa V, Kim HK, Missiakas DM, Schneewind O. Staphylococcal manipulation of host immune responses. Nat Rev Microbiol. 2015;13:529–43.

[44] Boyle-Vavra S, Li X, Alam MT, Read TD, Sieth J, Cywes-Bentley C, et al. USA300 and USA500 clonal lineages of *Staphylococcus aureus* do not produce a capsular polysaccharide due to conserved mutations in the *cap5* locus. MBio. 2015;6:e02585–14.

[45] Fattom AI, Horwith G, Fuller S, Propst M, Naso R. Development of StaphVAX, a polysaccharide conjugate vaccine against *S. aureus* infection: from the lab bench to phase III clinical trials. Vaccine. 2004;22:880–7.

[46] Thomer L, Schneewind O, Missiakas D. Pathogenesis of *Staphylococcus aureus* Bloodstream Infections. Annu Rev Pathol. 2016;11:343–64.

