## Supplemental Information for "A Protein A based *Staphylococcus aureus* vaccine with improved safety"

[†] Deceased 26 May 2019

**Key words**: *Staphylococcus aureus*; vaccine; protein A; anaphylaxis; IgE; B cell; superantigen; bloodstream infection; colonization; safety

**Conflict of interest**. The authors declare a competing financial interest as the inventors of patents describing SpA variants as vaccine against *S. aureus*. DM is the founder of ImmunArtes LLC, a University of Chicago startup company that aim to develop vaccines and therapies against *S. aureus* infections.

**Contributions.** M.S., X.C., H.K.K., O.S., and D.M. were involved in the design of experiments, analyses, and interpretation of data. M.S., X.C., Y.S., and O.S. performed experiments. O.S. and D.M. prepared the manuscript. All authors provided feedback and assisted in the editing of the manuscript.

**The Supplemental information contains:**

**Supplementary methods**
**Supplemental Figure 1**
**Supplemental Tables 1-4**

### Supplementary methods

#### Bacterial strains and growth conditions

*S. aureus* strains USA300 (LAC) and WU1 were grown in tryptic soy broth (TSB) or tryptic soy agar (TSA) at 37°C. Of note, isolate WU1 used in this study is resistant to rifampin which facilitates colony enumeration on TSA containing 100 µg/ml rifampin. *Escherichia coli* strains DH5α and BL21(DE3) were grown at 37°C in lysogeny broth (LB) medium with 100 µg/ml ampicillin and 1 mM isopropyl β-d-1-thiogalactopyranoside (IPTG) for the production of recombinant proteins.

#### Construction of SpA variants

The coding sequence of SpA variants was synthesized by Integrated DNA Technologies, Inc. The sequences and plasmid pET15b+ were digested by *Nde*I and *Bam*HI, respectively. The two digested products were ligated and transformed into *Escherichia coli* DH5α to generate the clones expressing N-terminal hexahistidine (His6)-tagged recombinant proteins. Candidate clones were validated by DNA sequencing. The correct plasmids were transformed into *E. coli* BL21 (DE3) for production of SpA variant candidates.

#### Purification of proteins

Cultures of *E. coli* (2 liters) that had been grown in LB supplemented with ampicillin and IPTG to an absorbance at 600 nm ($A_{600}$) of 2.0 were centrifuged (10,000 × *g* for 10 minutes). Sedimented cells were suspended in *Buffer A* (50 mM Tris-HCl [pH 7.5], 150 mM NaCl), and the resulting suspensions were lysed in a French press at 14,000 lb/in$^2$ (Thermo Spectronic, Rochester, NY). Unbroken cells were removed by centrifugation (5,000 × *g* for 15 minutes), and the crude lysates subjected to ultracentrifugation (100,000 × *g* for 1 hour at 4°C). Soluble recombinant proteins were subjected via gravity flow to chromatography on Ni-NTA agarose

(QIAGEN) with a packed volume of 1 ml preequilibrated with *Buffer A*. The columns were washed with 20 bed volumes of *Buffer A*, 20 bed volumes of *Buffer A* containing 10 mM imidazole, and eluted with 6 ml of *Buffer A* containing 500 mM imidazole. Aliquots of the eluted fractions were mixed with equal volumes of sample buffer and separated on 15% sodium dodecyl sulfate-polyacrylamide gel electrophoresis (SDS-PAGE) gels. Recombinant proteins were dialyzed against phosphate-buffered saline (PBS) and their concentrations determined with the bicinchoninic acid assay (Pierce). For immunization and for incubation with cell lines, recombinant protein preparations were subjected to the Endotoxin Removal Spin Columns (Pierce) to eliminate contaminating LPS. Sample purity was tested with ToxinSensor™ Chromogenic LAL Endotoxin Assay Kit (Genscript).

**Purification of antibodies**

To purify $V_H3$ IgG, human plasma (20 ml) prepared using whole human blood was subjected to affinity chromatography over Protein G Resin (Genscript) in order to remove human IgM, IgD, and IgA. Immunoglobulins eluted from Protein G Resin were subjected to a second affinity chromatography, $SpA_{KK}$-coupled resin, to enrich for $V_H3$ IgG [$SpA_{KK}$ cannot bind the Fcγ domain of IgG [1]]. Protein G Resin and $SpA_{KK}$-coupled resin were washed with 20-column volumes of PBS and bound proteins eluted with 0.1M glycine pH 3.0, neutralized with 1 M Tris-HCl, pH 8.5, and dialyzed against PBS overnight. For $V_H3$ IgE purification, the human cell line HEK 293F was used for transient expression of pVITRO1-Transtuzumab-IgE-κ. Cells were grown in DMEM/HIGH GLUCOSE medium with 10% FCS, 2 mM glutamine, penicillin (5,000 U/ml), and streptomycin (100 μg/ml). Cells transfected with pVITRO1-Transtuzumab-IgE-κ using PEI were incubated at 37°C in a 5% $CO_2$ atmosphere. For the stable expression of IgE, cells were cultured in Freestyle 293 medium for 7 days, and harvested at 12000 x *g* for 20 minutes. The supernatant was purified over 2 ml Protein L Resin (Genscript). The resin was washed with 20-column volumes of PBS, and bound $V_H3$ IgE was eluted and dialyzed as described above.

**Surface Plasmon Resonance (SPR)**

SPR experiments were performed on ProteOn™ XPR36 with ProteOn HTG chip. The running buffer was PBS with 0.05% Tween-20. The sensor-chip surfaces were activated with 2 mM nickel sulfate and regenerated with 300 mM EDTA, respectively. 500 nM of test articles (SpA wild type or variants) were immobilized at a flow rate of 25 µl/min. To measure interactions with wild type SpA, ligands (purified immunoglobulins) were used at concentrations of 500, 400, 300, 200 and 100 nM. To measure interactions with SpA variants, ligands were used at concentrations of 4, 3, 2, 1 and 0.5 µM. The association and dissociation rates were measured at a continuous flow rate of 30 µl/min and analyzed using the two-state reaction model. Association constants were determined from three independent experiments.

**Bio-layer Interferometry (BLI)**

The BLI experiment was performed using BLItz Bio-Layer Interferometer. Test candidates (25-50 nM) were immobilized onto a Ni-NTA sensor for 120 seconds. The sensor was equilibrated with PBS for 80 seconds, then dipped in solutions containing the ligand at concentrations of 20, 15, 10, and 0 µM for 120 seconds (association phase), followed by 120 seconds in PBS (dissociation phase). The data was acquired using BLI Data acquisition software 9.0 (FortéBIO) and analyzed using the Data Analysis software 9.0.0.14 (FortéBIO). Curve fitting was used to derive values of association constants.

**Enzyme-Linked Immunosorbent Assay (ELISA)**

Microtiter plates (NUNC MaxiSorp) were coated with purified antigens at 1 µg/ml (to measure antibody titers in test sera) or at 0.5 µg/ml (to measure interaction with 3F6 antibodies) in 0.1 M carbonate buffer (pH 9.5) at 4°C overnight. Wells were blocked, incubated with test serum or 3F6 monoclonal antibody prior to incubation with 1 mg/ml horseradish peroxidase (HRP)-conjugated mouse (Fisher Scientific) or (HRP)-conjugated human IgG (Jackson ImmunoResearch), and developed using OptEIA reagent (BD Biosciences). Half max titers were calculated with the GraphPad Prism software. The association constant was calculated from nonlinear regression (curve fit) model in the GraphPad Prism software. All experiments were performed in triplicate to calculate averages and standard error of the mean, and repeated for reproducibility.

**Anaphylactic response in µMT mice**

Mice with the µMT mutation were purchased from the Jackson Laboratory and bred at the University of Chicago. Cohorts of 4 six-week old female mice per group were sensitized by intradermal injection in the ear with human $V_H3$ IgG (2 µg in 20 µl of PBS) and 24 hours later, injected intravenously under anesthesia with ketamine–xylazine (100 mg–20 mg/kg) into the periorbital venous sinus of the right eye, with either PBS, SpA or its variants (200 µg in 100 µl PBS). Following a 5 minutes stimulation with the test article, animals were injected intravenously into the periorbital venous sinus of the left eye with 100 µl of 2% Evans blue. Animals were killed, their ears were dissected, dried, and extracted in formamide for 24 hours at 65°C. Evans blue extravasation in ear tissues (vascular permeability) was quantified by measuring absorbance at 620 nm. The experiment was performed twice. A representative experiment is shown.

**Human basophil activation experiments**

Blood (10 ml) was obtained from healthy donors and immediately mixed with 1 ml EDTA 0.1 M, pH7.5. SpA wild-type, vaccine candidate variants (1 μg), or PBS were added to 1-ml EDTA blood aliquots and the samples were incubated for 1 hour at 37°C with rotation. Sample aliquots were treated with RBC lysis buffer (Biolegend), centrifuged (350 x *g*) and supernatants discarded. Cells in pellets were washed in cold PBS and re-suspended in PBS with 5% FBS for staining with anti-CD123-FITC, anti-HLA-DA-PerCP, anti-CD63-PE, and anti-CD203c-APC (Biolegend) in the dark at room temperature for 10 min as described [2]. All stained samples were analyzed using a flow cytometer (BD LSRII 3-8, BD Biosciences). Total basophil counts were obtained by gating from $SSC^{low}$/$CD203c^{+}$/$CD123^{+}$/$HLA\text{-}DR^{-}$ cells and activated basophils were selected from the $CD63^{+}CD203c^{+}$ pool. Experiments were performed in triplicate and repeated at least three times using different healthy donors.

**Mast cell degranulation**

Human mast cells (LAD2) [kindly provided by Dr. Kirshenbaum from NIAID] were sensitized by incubating 2 x $10^5$ cells with 100 ng human $V_H3$ IgE overnight at 37°C in a 5% $CO_2$ atmosphere. Cells were harvested and washed twice with HEPES buffer containing 0.04% bovine serum albumin (BSA) to remove free IgE. Cells were suspended in the same buffer at the concentration of 2 x $10^5$ cells/ml, and stimulated with SpA or test articles for 30 min before assaying for β-hexosaminidase and histamine release. Cells were sedimented and the spent medium was transferred to a fresh tube while cells in the pellet were lysed with 0.1% Triton X-100. β-hexosaminidase activity in the spent medium and in the Triton X-100-lysed cells, was measured by adding the colorimetric substrate pNAG (p-nitrophenyl-N-acetyl-β-D-glucosaminide obtained from Sigma; final concentration 3.5 mg/ml at pH 4.5) for 90 min. The reaction was quenched by the addition of 0.4 M glycine pH 10.7 and the absorbance recorded at

405 nm. The results were expressed as the percentage of β-hexosaminidase released in the spent medium over total (spent medium + Triton X-100-lysed cells). Experiments were performed in triplicate and repeated at least three times. Histamine was measured using an Enzyme Immunoassay (SpiBio Bertin Pharma). Briefly, wells of a microtiter plate were coated with mouse anti-histamine antibody and incubated for 24 hours with tracer (acetylcholinesterase linked to histamine) mixed with an experimental extract. Plates were washed, and Ellman's Reagent (acetylcholinesterase substrate) was added to the wells. Product formation was detected by recording absorbance at 412 nm. Absorbance at 412 nm is proportional to the amount of tracer bound to the well and is inversely proportional to the amount of histamine present in the experimental extract. All samples were performed in duplicate.

**Active immunization of mice**

Animals BALB/c or C57BL/6J (3-week-old, female mice, 10-15 animals per group) were immunized with PBS, or 50 µg purified endotoxin-free protein $SpA_{KKAA}$ or $SpA_{Q9,10K/S33E}$ or $SpA_{Q9,10K/S33T}$ emulsified in 5:2:3 of antigen: CFA: IFA and boosted with 50 µg proteins emulsified in 1:1 of antigen: IFA 11 days following the first immunization. On day 20, mice were bled and sera were harvested to evaluate antibody titers to vaccine candidates by ELISA. On day 21, mice were either inoculated for nasopharyngeal colonization or infected by the intravenous injection of bacteria.

**Mouse nasopharyngeal colonization**

Overnight cultures of the *S. aureus* strain WU1 were diluted 1:100 in fresh TSB and grown for 2 h at 37°C as described [3]. The cells were centrifuged, washed, and suspended in PBS. C57BL/6J mice (Jackson Laboratory) were anesthetized by intraperitoneal injection with ketamine–xylazine (100 mg–20 mg/kg), and 1 x $10^8$ CFU of *S. aureus* (in a 10-µl volume) was pipetted into the right nostril of each mouse. Inocula were quantified by spreading sample aliquots on TSA and enumerating the colonies that formed upon incubation. In weekly intervals following inoculation, the oropharynx of the mice was swabbed and stool samples were collected and homogenized in PBS. Swab samples and homogenates of stool samples were spread on TSA containing 100 μg/ml rifampin for bacterial enumeration, a process that effectively restricted growth of the normal flora. At the end of the experiment, the mice were bled via periorbital vein puncture to obtain sera for antibody response analyses using the staphylococcal antigen matrix as described [4]. Briefly, nitrocellulose membranes were blotted with 2 μg affinity-purified staphylococcal antigens. The membranes were blocked with 5% degranulated milk and incubated with diluted mouse sera (1:10,000 dilution) and IRDye 680-conjugated goat anti-mouse IgG (Li-Cor). Signal intensities were quantified using the Odyssey infrared imaging system (Li-Cor). Nasopharyngeal and stool colonization, ELISA, and antigen matrix experiments were performed at least twice.

**Mouse renal abscess model**

Inocula of *S. aureus* USA300 (LAC) were prepared as described for strain WU1. Bacterial suspensions ($5 \times 10^6$ CFU) were inoculated into the periorbital venous sinus of the right eye of anesthetized BALB/c mice. On day 15, following challenge, mice were killed by $CO_2$ inhalation. Both kidneys were removed, and the staphylococcal load in one organ was analyzed by homogenizing renal tissue with PBS, 0.1% Triton X-100. Serial dilutions of homogenate were spread on TSA and incubated for colony formation. The remaining organ was examined by histopathology. Briefly, kidneys were fixed in 10% formalin for 24 h at room temperature. Tissues were embedded in paraffin, thin sectioned, stained with hematoxylin-eosin, and inspected by light microscopy to enumerate abscess lesions. The experiment was performed twice.

### REFERENCES


[1] Falugi F, Kim HK, Missiakas DM, Schneewind O. The role of protein A in the evasion of host adaptive immune responses by *Staphylococcus aureus* mBio. 2013;4:e00575-13.

[2] Santos AF, Becares N, Stephens A, Turcanu V, Lack G. The expression of CD123 can decrease with basophil activation: implications for the gating strategy of the basophil activation test. Clin Transl Allergy. 2016;6:11.

[3] Sun Y, Emolo CE, Holtfreter S, Wiles S, Kreiswirth B, Missiakas D, et al. Staphylococcal protein A is required for persistent colonization of mice with *Staphylococcus aureus*. J Bacteriol. 2018;200:e00735-17.

[4] Kim HK, Cheng AG, Kim H-Y, Missiakas DM, Schneewind O. Non-toxigenic protein A vaccine for methicillin-resistant *Staphylococcus aureus* infections. J Exp Med. 2010;207:1863-70.

**Supplementary Figure 1**

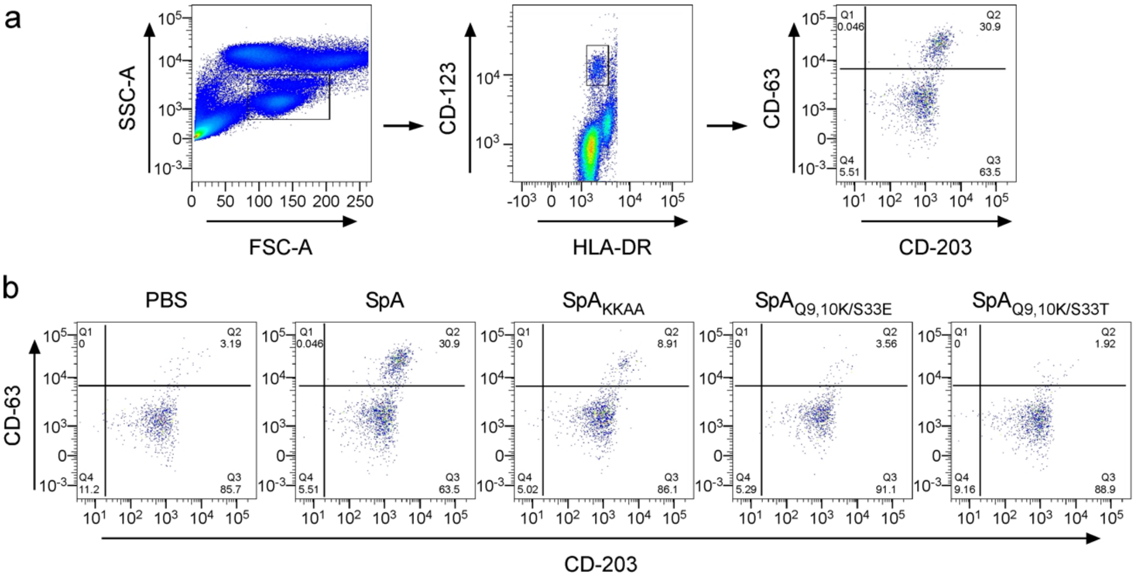


**Supplementary Fig. 1. Activation of human basophils by vaccine candidate variants. a** | A stepwise process was used to sort and enumerate activated basophil after incubation of EDTA-treated human blood with wild-type SpA as described [2]. The three sorting steps are shown. **b** | Representative images for the final sorting step of activated human basophils are shown for PBS, SpA (same image as in panel **a**), and vaccine variants: $SpA_{KKAA}$, $SpA_{Q9,10K/S33E}$, $SpA_{Q9,10K/S33T}$. Refer to Table 3 for the quantification of the data.

### Supplementary Tables 1-4

**Supplementary Table 1. Affinity measurements with wild-type SpA, $SpA_{KKAA}$ and $SpA_{Q9,10K/G29X}$ vaccine candidates and human IgG[#].**

| $SpA_{Q9,10K/G29X}$[a] | $K_A$ ($\times 10^5$ $M^{-1}$)[b] | SD ($\times 10^5$)[c] | *P* value[d] |
|---|---|---|---|
| $SpA_{Q9,10K/G29H}$ | 1.435 | 0.2799 | * |
| $SpA_{Q9,10K/G29C}$ | 1.743 | 0.8619 | * |
| $SpA_{Q9,10K/G29T}$ | 1.982 | 0.9146 | ns |
| $SpA_{Q9,10K/G29Q}$ | 2.057 | 0.9600 | * |
| $SpA_{Q9,10K/G29L}$ | 3.146 | 1.3860 | ns |
| $SpA_{Q9,10K/G29E}$ | 3.182 | 1.5300 | ns |
| $SpA_{Q9,10K/G29P}$ | 3.396 | 1.4410 | ns |
| $SpA_{Q9,10K/G29F}$ | 3.460 | 1.5860 | ns |
| $SpA_{Q9,10K/G29M}$ | 3.893 | 0.7868 | ns |
| $SpA_{Q9,10K/G29V}$ | 4.350 | 1.0830 | ns |
| $SpA_{Q9,10K/G29W}$ | 4.508 | 0.7448 | ns |
| $SpA_{Q9,10K/G29D}$ | 5.478 | 1.0150 | ns |
| $SpA_{Q9,10K/G29R}$ | 6.056 | 0.9814 | ns |
| $SpA_{Q9,10K/G29N}$ | 6.231 | 0.7696 | ns |
| $SpA_{Q9,10K/G29Y}$ | 8.367 | 3.326 | ns |
| $SpA_{Q9,10K/G29S}$ | 9.398 | 4.298 | *** |
| $SpA_{Q9,10K/G29K}$ | 9.738 | 2.345 | ** |
| $SpA_{Q9,10K/G29I}$ | 10.070 | 4.398 | ** |
| $SpA_{Q9,10K/G29A}$ | 11.310 | 3.119 | *** |
| **$SpA_{KKAA}$** | **5.022** | **2.150** | |
| SpA | 1081 | 16.34 | |

[a]Test articles were immobilized on Bio-Rad ProteOn HTG Chip and subjected to Surface Plasmon Resonance measurements with increasing concentrations of human IgG and flowed over each channel of the chip. Data were analyzed from three independent experimental determinations.
[b]Data were used to derive the association constant ($K_A$) for each test article.
[c]Data were used to derive the Standard Deviation (SD) for each test article.
[d]Data were analyzed with One-way ANOVA with Dunnett's Multiple Comparison Test between each test article and $SpA_{KKAA}$. Symbols: ns, not significant; *, $P<0.05$; **, $P<0.01$; ***, $P<0.001$; ****, $P<0.0001$.

**Supplementary Table 2. Affinity measurements with wild-type SpA, $SpA_{KKAA}$ and $SpA_{Q9,10K/S33X}$ vaccine candidates and human IgG[#].**

| **$SpA_{Q9,10K/S33X}$**[a] | ***$K_A$*** ($\times10^5$ $M^{-1}$)[b] | **SD** ($\times10^5$)[c] | ***P* value**[d] |
|---|---|---|---|
| $SpA_{Q9,10K/S33E}$ | 0.496 | 0.0439 | ** |
| $SpA_{Q9,10K/S33T}$ | 0.386 | 0.1218 | *** |
| $SpA_{Q9,10K/S33Y}$ | 1.571 | 0.7497 | ns |
| $SpA_{Q9,10K/S33YI}$ | 1.840 | 1.1290 | * |
| $SpA_{Q9,10K/S33L}$ | 2.051 | 0.7592 | ns |
| $SpA_{Q9,10K/S33W}$ | 2.356 | 0.6373 | ns |
| $SpA_{Q9,10K/S33V}$ | 2.471 | 1.2060 | ns |
| $SpA_{Q9,10K/S33H}$ | 2.784 | 0.6087 | ns |
| $SpA_{Q9,10K/S33N}$ | 3.066 | 1.0100 | ns |
| $SpA_{Q9,10K/S33M}$ | 3.177 | 1.3750 | ns |
| $SpA_{Q9,10K/S33R}$ | 3.463 | 1.7950 | ns |
| $SpA_{Q9,10K/S33D}$ | 3.824 | 1.7100 | ns |
| $SpA_{Q9,10K/S33F}$ | 3.902 | 1.8040 | ns |
| $SpA_{Q9,10K/S33Q}$ | 4.068 | 2.8350 | ns |
| $SpA_{Q9,10K/S33P}$ | 4.218 | 2.2560 | ns |
| $SpA_{Q9,10K/S33C}$ | 4.577 | 0.6927 | ns |
| $SpA_{Q9,10K/S33K}$ | 5.124 | 2.1810 | ns |
| $SpA_{Q9,10K/S33A}$ | 10.540 | 5.0520 | *** |
| $SpA_{Q9,10K/S33G}$ | 11.180 | 5.2040 | *** |
| **$SpA_{KKAA}$** | **5.022** | **0.0439** | |
| SpA | $1081^{-1}$ | 16.34 | |

[a]Test articles were immobilized on Bio-Rad ProteOn HTG Chip and subjected to Surface Plasmon Resonance measurements with increasing concentrations of human IgG and flowed over each channel of the chip. Data were analyzed from three independent experimental determinations.

[b]Data were used to derive the association constant ($K_A$) for each test article.

[c]Data were used to derive the Standard Deviation (SD) for each test article.

[d]Data were analyzed with One-way ANOVA with Dunnett's Multiple Comparison Test between each test article and $SpA_{KKAA}$. Symbols: ns, not significant; *, $P<0.05$; **, $P<0.01$; ***, $P<0.001$; ****, $P<0.0001$.

**Supplementary Table 3. The association constant for binding to human IgG of SpA variants Q9,10K/S33X or Q9,10K/G29X in combination with other amino acid substitutions[#]**

| **Parent SpA variant[a]** | **$K_A$ (x10$^5$ M$^{-1}$)[b]** | **SD[c]** | **$P$ value[d]** | **Parent SpA variant with additional substitutions[a]** | **$K_A$ (x10$^5$ M$^{-1}$)[b]** | **SD (x10$^5$ M$^{-1}$)[c]** | **$P$ value[e]** |
|---|---|---|---|---|---|---|---|
| SpA$_{Q9,10K/S33E}$ | 0.496 | 0.044 | ** | SpA$_{Q9,10K/S33E/D36,37A}$ | 0.568 | 0.1185 | ns |
| | | | | SpA$_{Q9,10K/S33E/G29A}$ | 1.265 | 0.6947 | ns |
| | | | | SpA$_{Q9,10K/S33E/D36,37A/G29A}$ | 1.892 | 0.6793 | ns |
| | | | | SpA$_{Q9,10K/S33E/G29F}$ | 1.575 | 0.4060 | ns |
| | | | | SpA$_{Q9,10K/S33E/D36,37A/G29F}$ | 14.850 | 13.480 | *** |
| | | | | SpA$_{Q9,10K/S33E/G29R}$ | 4.840 | 1.1960 | ns |
| | | | | SpA$_{Q9,10K/S33E/D36,37A/G29R}$ | 10.240 | 5.2600 | * |
| SpA$_{Q9,10K/S33Q}$ | 4.068 | 2.835 | ns | SpA$_{Q9,10K/S33Q/D36,37A}$ | 3.930 | 1.9290 | ns |
| | | | | SpA$_{Q9,10K/S33Q/G29A}$ | 2.563 | 1.3670 | ns |
| | | | | SpA$_{Q9,10K/S33Q/D36,37A/G29A}$ | 4.893 | 3.8360 | ns |
| | | | | SpA$_{Q9,10K/S33Q/G29F}$ | 1.275 | 0.7355 | ns |
| | | | | SpA$_{Q9,10K/S33Q/D36,37A/G29F}$ | 12.470 | 8.8810 | * |
| | | | | SpA$_{Q9,10K/S33Q/G29R}$ | 2.333 | 0.4245 | ns |
| | | | | SpA$_{Q9,10K/S33Q/D36,37A/G29R}$ | 6.378 | 4.6820 | ns |
| SpA$_{Q9,10K/S33F}$ | 3.902 | 1.804 | ns | SpA$_{Q9,10K/S33F/D36,37A}$ | 3.634 | 2.6420 | ns |
| | | | | SpA$_{Q9,10K/S33F/G29A}$ | 1.190 | 0.4299 | ns |
| | | | | SpA$_{Q9,10K/S33F/D36,37A/G29A}$ | insoluble | | |
| | | | | SpA$_{Q9,10K/S33F/G29F}$ | 2.440 | 0.7657 | ns |
| | | | | SpA$_{Q9,10K/S33F/D36,37A/G29F}$ | insoluble | | |
| | | | | SpA$_{Q9,10K/S33F/G29R}$ | 1.903 | 0.8693 | ns |
| | | | | SpA$_{Q9,10K/S33F/D36,37A/G29R}$ | 9.056 | 4.9730 | * |
| SpA$_{Q9,10K/S33K}$ | 5.124 | 2.181 | ns | SpA$_{Q9,10K/S33K\ /D36,37A}$ | 8.048 | 4.1050 | ns |
| SpA$_{Q9,10K/S33A}$ | 10.540 | 5.052 | *** | SpA$_{Q9,10K/S33A/D36,37A}$ | 18.830 | 18.320 | ns |
| SpA$_{Q9,10K/G29F}$ | 3.460 | 1.586 | ns | SpA$_{Q9,10K/G29F/D36,37A}$ | 3.723 | 1.5100 | ns |
| SpA$_{Q9,10K/G29R}$ | 6.056 | 0.981 | ns | SpA$_{Q9,10K/G29R/D36,37A}$ | 6.808 | 3.6840 | ns |
| SpA$_{Q9,10K/G29A}$ | 11.310 | 3.119 | *** | SpA$_{Q9,10K/G29A/D36,37A}$ | 1.78 | 0.5098 | *** |
| SpA-KR | 5.464 | 0.767 | ns | | | | |
| SpA$_{RRVV}$ | 5.609 | 2.355 | ns | | | | |
| **SpA$_{KKAA}$** | **5.022** | **2.150** | **-** | | | | |

[a]Test articles were immobilized on Bio-Rad ProteOn HTG Chip and subjected to Surface Plasmon Resonance measurements with increasing concentrations of human IgG and flowed over each Chip. Data were analyzed from three independent experimental determinations.
[b]Data were used to derive the association constant ($K_A$) for each test article.
[c]Data were used to derive the standard deviation (SD) for each test article.
[d,e]Data were analyzed with One-way ANOVA with Dunnett's Multiple Comparison Test between test article and SpA$_{KKAA}$[d] and between test article (column 5) and parent vaccine (column 1)[e]. Symbols: ns, not significant; *, $P$<0.05; **, $P$<0.01; ***, $P$<0.001; ****, $P$<0.0001.

**Supplementary Table 4. Impact of immunization with SpA variants on serum IgG responses in *S. aureus* WU1-colonized C57BL/6J mice**

| | SpA$_{KKAA}$ immunized | | | | SpA$_{Q9,10K/S33E}$ immunized | | | | SpA$_{Q9,10K/S33T}$ immunized | | | | PBS immunized | |
|---|---|---|---|---|---|---|---|---|---|---|---|---|---|---|
| | Colonized | | Cleared | | colonized | | cleared | | colonized | | cleared | | colonized | |
| | Fold change | *P*[1] | Fold change | *P*[2] | Fold change | *P*[1] | Fold change | *P*[3] | Fold change | *P*[1] | Fold change | *P*[4] | Fold change | *P*[1] |
| SpA$_{KKAA}$ | 33.87±2.860 | **** | 30.02±8.985 | ns | 33.34±14.86 | **** | 35.9±5.004 | ns | 19.49±6.939 | * | 37.88±14.33 | ns | 1.34±1.174 | ns |
| ClfA | 4.18±2.269 | ns | 6.51±2.530 | ns | 4.54±2.585 | * | 5.09±0.350 | ns | 1.26±0.292 | ns | 5.577±1.739 | * | 1.12±1.145 | ns |
| ClfB | 1.88±3.433 | ns | **33.67±35.60** | * | 4.77±4.526 | ns | 39.49±30.69 | ** | 2.17±2.837 | ns | 2.09±0.728 | ns | 0.67±1.237 | ns |
| SdrC | 8.81±1.737 | ns | 17.21±10.590 | ns | 18.82±11.35 | ns | 23.43±13.23 | ns | 6.96±2.248 | ns | 11.89±4.715 | ns | 8.02±1.854 | ns |
| SdrD | 6.35±2.830 | ns | **22.75±17.030** | *** | 2.41±1.637 | ns | **15.55±8.702** | *** | 3.17±3.459 | ns | 1.53±1.444 | ns | 1.28±1.521 | ns |
| SdrE | 3.97±3.192 | ns | 33.92±31.170 | ns | 13.72±13.23 | ns | **64.73±45.82** | ** | 5.54±3.062 | ns | 2.77±5.791 | ns | 1.38±2.433 | ns |
| SasI | 5.34±1.088 | * | **14.36±1.320** | **** | 5.94±2.638 | ** | 10.08±4.768 | * | 3.18±0.530 | ns | 4.86±1.051 | ns | 1.45±0.739 | ns |
| Coa | 5.34±3.239 | ns | 0.63±1.085 | ns | 2.58±2.057 | ns | 4.84±4.277 | ns | 1.25±1.097 | ns | 4.31±5.357 | ns | 3.31±2.459 | ns |
| Hla | 3.10±1.187 | ns | 2.92±0.798 | ns | 1.37±0.549 | ns | 1.63±1.635 | ns | 0.97±0.450 | ns | 1.50±0.730 | ns | 1.98±0.460 | ns |
| SasF | 1.75±1.491 | ns | 1.83±3.175 | ns | 16.59±12.34 | ns | 16.22±6.612 | ns | 3.64±2.847 | ns | 17.26±19.98 | ns | 3.45±6.995 | ns |
| SasK | 4.99±1.971 | ns | 6.91±6.150 | ns | 1.18±1.02 | ns | 4.3±1.574 | ns | 0.60±0.393 | ns | 1.06±1.204 | ns | 2.17±1.244 | ns |
| SasD | 2.86±2.619 | ns | 4.06±3.675 | ns | 1.68±0.831 | ns | **8.98±1.789** | **** | 0.83±0.187 | ns | 1.88±1.155 | ns | 1.41±0.844 | ns |
| SasA | 7.83±3.352 | ns | **20.70±4.964** | *** | 8.92±3.843 | ns | **21.08±2.179** | **** | 4.29±0.473 | ns | 5.65±4.237 | ns | 5.00±1.902 | ns |
| IsdA | 1.27±0.549 | ns | 1.61±0.503 | ns | 1.46±0.268 | ns | 1.58±0.282 | ns | 1.21±0.156 | ns | 1.72±0.3081 | ns | 1.47±0.427 | ns |
| IsdB | 5.37±2.493 | ns | 18.26±6.296 | ns | 14.52±11.03 | ** | 10.55±8.05 | ns | 2.26±1.103 | ns | 10.4±3.17 | ns | 2.50±2.509 | ns |
| FnbpA | 2.93±0.720 | ns | **8.60±7.333** | * | 4.01±2.275 | ns | **13.4±2.257** | **** | 2.12±0.593 | ns | 2.54±1.839 | ns | 1.63±1.130 | ns |
| FnbpB | 5.72±4.609 | ns | **48.49±48.080** | ** | 17.63±11.82 | ns | **60.76±19.73** | *** | 10.53±1.502 | ns | 7.9±5.119 | ns | 5.12±5.783 | ns |
| SasG | 7.99±4.711 | * | **19.32±3.083** | ** | 9.24±6.548 | ** | **27.85±4.588** | **** | 10.08±0.942 | * | 9.00±7.45 | ns | 0.95±0.641 | ns |
| EsxB | 18.55±12.860 | ** | 24.38±10.140 | ns | 0.17±0.408 | ns | **22.57±17.27** | ** | no signal | / | 1.23±2.113 | / | 1.88±3.371 | ns |
| SCIN | 3.99±1.502 | * | 3.79±0.644 | ns | 0.79±0.851 | ns | **4.14±1.709** | ** | 0.41±0.290 | ns | 0.54±0.466 | ns | 1.91±1.193 | ns |
| Eap | 6.22±4.380 | * | 9.20±4.295 | ns | no signal | / | 0.08±0.119 | / | no signal | / | no signal | / | 2.38±0.904 | ns |
| vWbp | 14.06±9.443 | ns | 23.32±12.650 | ns | 5.29±4.661 | ns | **24.59±16.34** | ** | 4.45±1.663 | ns | 3.62±3.776 | ns | 2.02±4.291 | ns |
| Efb | 3.65±2.128 | ns | 7.06±3.743 | ns | 0.94±1.159 | ns | **7.92±6.77** | ** | 0.29±0.264 | ns | 0.19±0.329 | ns | 2.10±0.611 | ns |
| LukS | 0.72±1.247 | ns | no signal | / | 10.70±13.22 | * | 3.47±4.624 | ns | 5.11±3.76 | ns | 6.36±7.309 | ns | 0.64±0.700 | ns |
| LukF | 0.88±1.760 | ns | 1.36±2.107 | ns | 14.14±8.156 | *** | 20.9±12.45 | ns | 16.34±5.141 | ** | 11±4.966 | ns | 0.81±1.168 | ns |

Fold changes were calculated by dividing the average signal intensities derived from *S. aureus*-inoculated mice by the average signal intensities from naïve mice
*P* values were calculated using one-way ANOVA with Sidak's multiple comparisons test.
The data are presented as means ± standard deviations. Bold values point to antibody titers significantly increased in decolonized animals.
Symbol "/": the signal was not detectable; ns: not significant; P<0.05; **, P<0.01; ***, P<0.001; ****, P<0.0001
$P^1$, $P^2$, $P^3$, $P^4$, Fold changes of average signal intensities were compared to: $P^1$, naïve mice; $P^2$, mice immunized with $SpA_{KKAA}$ but still colonized; $P^3$, mice immunized with $SpA_{Q9,10K/S33E}$ but still colonized; $P^4$, mice immunized with $SpA_{Q9,10K/S33T}$ but still colonized.
